# Activation induces shift in nutrient utilization that differentially impacts cell functions in human neutrophils

**DOI:** 10.1101/2023.09.25.559385

**Authors:** Emily C. Britt, Xin Qing, James A. Votava, Jorgo Lika, Andrew Wagner, Simone Shen, Nicholas L. Arp, Hamidullah Khan, Stefan M. Schieke, Christopher D. Fletcher, Anna Huttenlocher, Jing Fan

## Abstract

Neutrophils – the first responders in innate immunity – perform a variety of effector functions associated with specific metabolic demand. To maintain fitness and support functions, neutrophils have been found to utilize extracellular glucose, intracellular glycogen, and other alternative substrates. However, the quantitative contribution of these nutrients under specific conditions and the relative dependence of various cell functions on specific nutrients remain unclear. Here, using *ex vivo* and *in vivo* isotopic tracing, we reveal that under resting condition, human peripheral blood neutrophils, in contrast to *in vitro* cultured human neutrophil-like cell lines, rely on glycogen as a major direct source of glycolysis and pentose phosphate pathway. Upon activation with a diversity of stimuli, neutrophils undergo a significant and often rapid nutrient preference shift, with glucose becoming the dominant metabolic source thanks to a multi-fold increase in glucose uptake mechanistically mediated by the phosphorylation and translocation of GLUT1. At the same time, cycling between gross glycogenesis and glycogenolysis is also substantially increased, while the net flux favors sustained or increased glycogen storage. The shift in nutrient utilization impacts neutrophil functions in a function-specific manner. The activation of oxidative burst specifically depends on the utilization of extracellular glucose rather than glycogen. In contrast, the release of neutrophil traps can be flexibly supported by either glucose or glycogen. Neutrophil migration and fungal control is promoted by the shift away from glycogen utilization. Together, these results quantitatively characterize fundamental features of neutrophil metabolism and elucidate how metabolic remodeling shapes neutrophil functions upon activation.

## Introduction

Neutrophils, the most abundant circulating leukocytes, provide the first line of defense in innate immunity. In this role, neutrophils are charged with metabolically demanding immune functions such as oxidative burst, phagocytosis, and neutrophil extracellular trap (NET) release^1,2^. Glycolysis has long been accepted as the main source of energy to support mature neutrophils’ dynamically changing metabolic demands^3^, while neutrophil mitochondria has limited contribution^4,5^. The pentose phosphate pathway (PPP) has also been found critical for some immune functions in neutrophils, particularly oxidative burst^6–9^.

These crucial metabolic pathways can be supplied by different nutrient sources, including exogenous glucose or intracellular glycogen storage, both of which feed into the production of glucose-6-phosphate (G6P), the common precursor for glycolysis and PPP activity. Prior research has suggested the importance of both glucose metabolism and glycogenolysis in immunity. For instance, increased glucose uptake by GLUT1 is essential for CD4 T cell activation and function^10^, while glycogenolysis is essential for fueling CD8 memory T cell recall and the early response of dendritic cell activation^11,12^. Particularly in neutrophils, glucose has been traditionally considered as the main nutrient source, and uptake of extracellular glucose has been reported to increase upon exposure to classical activators or pathogens (including *Francisella tularensis, leishmania,* and *Candida albicans*)^13–20^. On the other hand, recent studies have revealed neutrophils also possess meaningful glycogen storage, and glycogenolysis is increased by certain stimulations or diseases^3,5^. Moreover, despite neutrophils’ relatively few mitochondria and low mitochondrial energy generation, it has been reported that mitochondrial substrates can support some effector functions such as prolonged oxidative burst^21,22^, and that neutrophils express key gluconeogenic enzymes including mitochondrial phosphoenolpyruvate carboxykinase^5^, indicating that neutrophils have the potential to generate glycolytic intermediates or glycogen storage from mitochondrial gluconeogenic substrates.

These recent studies clearly suggest that neutrophils have the capacity to utilize a range of nutrients to support their metabolism, survival, and function, and that neutrophil activation is associated with significant reprogramming of nutrient metabolism. However, two fundamental questions remain unclear. First, what are the quantitative contributions of these implicated nutrients under specific conditions? Understanding the relative contribution would bring a complete picture of neutrophils’ *nutrient preference* in different states. Second, what is the *nutrient dependence* of various neutrophil functions? As the first innate immune cells to be recruited to sites of infection and injury, neutrophils can encounter changing microenvironments with varied nutrient levels. Some metabolic sources, though preferred under optimal conditions, may be compensated for by other options when unavailable, while others may be indispensable for certain effector functions. Determining (a) the extent to which neutrophils have the flexibility to use different metabolic sources to support the initiation and sustentation of their critical functions and (b) whether altering neutrophils’ nutrient utilization can steer their functions has important implications in health and disease.

Here we quantified the contribution of major metabolic sources in neutrophils in the resting state as well as upon activation by a variety of stimuli. In light of accumulating evidence suggesting that physiological environment and specific cell-intrinsic factors can significantly alter metabolism, we quantitatively evaluated the nutrient preference of human neutrophils in three setting: *in vivo* by isotopic tracing in healthy volunteers, *ex vivo* by immediately assessing freshly insolated human peripheral blood neutrophils in culture condition, and *in vitro* in a widely used stable neutrophil-like cell line cultured in media. This study reveals fundamental features of human neutrophil metabolism and elucidates the specific metabolic rewiring induced by different stimulations and conditions. We further quantitatively assessed the dependence of an array of important functions on the two major nutrient sources, extracellular glucose and intracellular glycogen, thus revealing the differential effects of metabolic perturbations, which mimic or reverse the stimulation-induced nutrient preference shift, on neutrophil functions. These findings have significant health relevance given that dysregulation of neutrophil functions contributes to many infectious diseases and inflammatory conditions and understanding the specific metabolism–function connections carries broad potential therapeutic applications.

## Results

### Primary human neutrophils use glycogen storage as a major direct metabolic source in the resting state

To quantitatively characterize human neutrophils’ nutrient preference under baseline condition, we started with evaluating the contribution of extracellular glucose, which has been traditionally recognized as the main source of neutrophil metabolism. Recent studies have highlighted that cellular metabolism can be significantly impacted by cell intrinsic factors as well as nutrient availability and extracellular signals in their environment^23,24^, causing knowledge gained using *in vitro* or animal models may not always translating into human contexts. Thus, we applied a cutting edge *in vivo* isotopic tracing approach to get a precise picture of human neutrophil metabolism in their physiological environment. Healthy volunteers were intravenously infused with uniformly labeled glucose tracer (U-^13^C glucose in sterile saline, infused with an initial bolus of 8g in 10 min, followed by a steady infusion rate of 8g per hour, following previously developed methods^25–27)^, and blood samples were collected at various time points for analysis of metabolites in plasma and in peripheral blood neutrophils (Fig. 1a).

**Figure 1.**
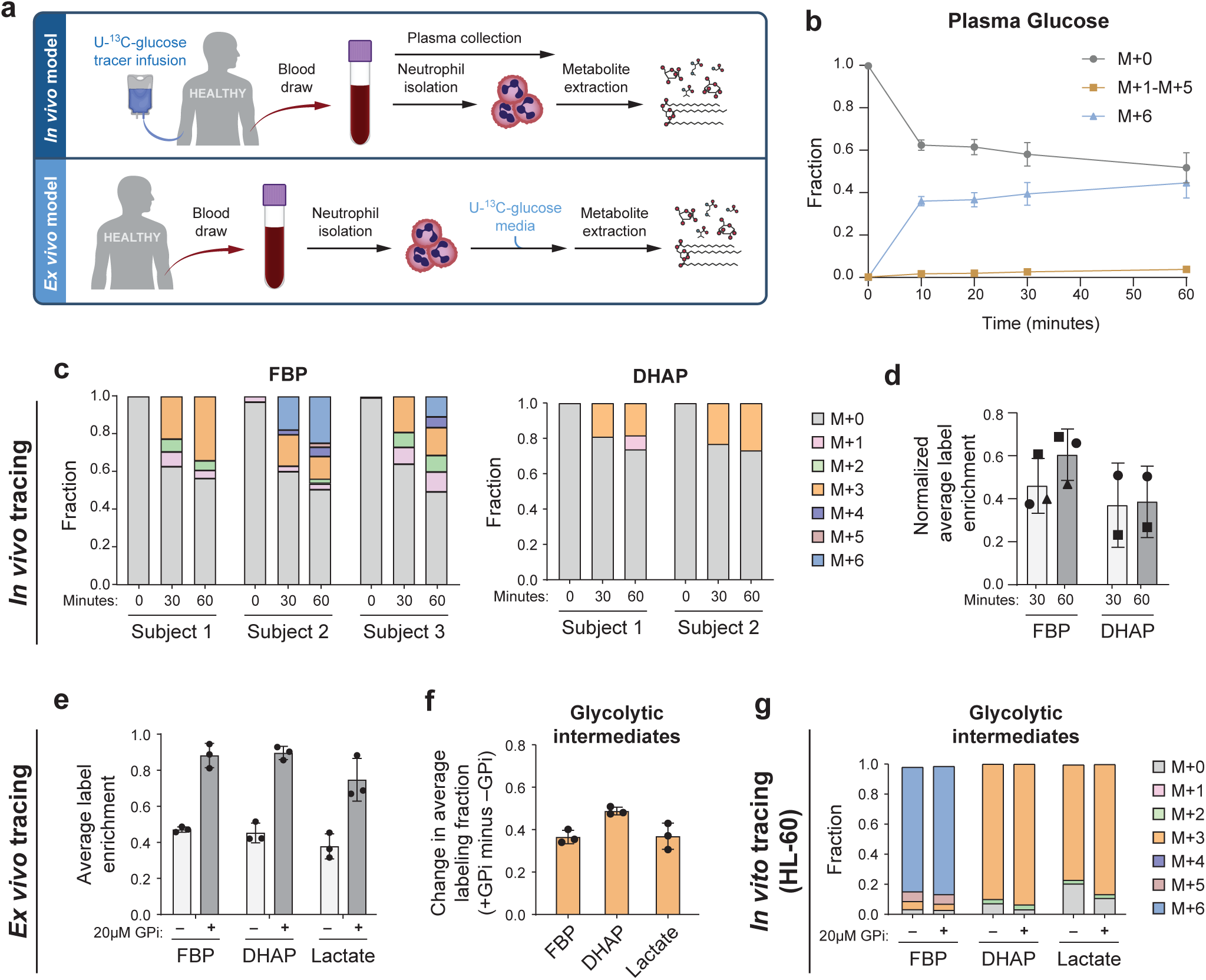
**Contribution of glucose and glycogen in the metabolism of human neutrophils in the resting state.** a. Schematic of *in vivo* and *ex vivo* U-^13^C-glucose tracing. b. Plasma glucose labeling over a time course following U-^13^C-glucose infusion. Results show mean ± SD from n=3 independent participants. c. Full isotopic labeling distribution of intracellular fructose-1,6-bisphosphate (FBP), and dihydroxyacetone phosphate (DHAP) from neutrophils isolated from blood sample pre-infusion (0 min), 30 or 60 min of U-^13^C-glucose infusion for each participant (DHAP signal was below limit for reliable quantification in subject 3). d. Average labeling enrichment (on a per carbon basis) of intracellular glycolytic metabolites from neutrophils isolated from blood after 30 or 60 min of U-^13^C-glucose infusion, normalized to the labeling enrichment of plasma glucose. Bars and error bars show mean ± SD from n=3 independent participants, represented by different markers. e. Average labeling enrichment of glycolytic metabolites (on a per carbon basis) from peripheral blood neutrophils incubated *ex vivo* in U-^13^C-glucose media for 30 min with or without treatment of 20μM glycogen phosphorylase inhibitor (GPi). Bars and error bars show mean ± SD from n=3 independent donors. f. Difference between labeling enrichment of glycolytic intermediates from unstimulated neutrophils treated with GPi compared to untreated. Bars and error bars show mean ± SD from n=3 independent donors. g. Full isotopic labeling distribution of intracellular glycolytic intermediates from HL-60 cells following 30-min incubation in U-^13^C-glucose media with or without GPi treatment. Results show the average of technical replicates from one representative experiment. Unstimulated labeling results were confirmed in three independent experiments.

In plasma, glucose labeling quickly enriched to ∼40% labeled within 10 min and stayed mostly steady up to 60 min (Fig. 1b), while total glucose level maintained a normal range (Supplementary Fig. 1a). During this time, most labeled glucose remained uniformly labeled, with a small fraction (<5%) of other label forms (M+1 to M+5) slowly arising over time, indicating metabolic turnover of glucose by tissues via gluconeogenesis. Plasma lactate, which has been recently identified as a major glucose-derived circulating metabolite^28,29^, gradually increased to ∼15% labeled (Supplementary Fig. 1b). Other circulating metabolites, including glycerol, which may directly contribute to lower glycolysis, glutamine, glutamate, and alanine, remained minimally labeled in plasma (Supplementary Fig. 1c–f). These are the major circulating metabolites available to neutrophils that can potentially deliver labeling in cells.

In neutrophils, intracellular glycolytic intermediates quickly gained labeling after infusion. The dominant labeling forms, including 3- and 6- labeled fructose-1,6-bisphosphate (FBP) and 3- labeled dihydroxyacetone phosphate (DHAP), reflect direct contribution of circulating glucose via glycolysis (Fig. 1c). A smaller fraction of other label forms, including 1- or 2- labeled, may have resulted from either scrambling non-oxidative PPP, which we found is highly reversible in resting neutrophils^6^, or to a lesser extent, from gross flux from phosphoenolpyruvate derived from phosphoenolpyruvate carboxykinase. There was a small fraction of labeling in intracellular TCA cycle intermediates during this time frame (Supplementary Fig. 1g). The average labeling enrichments of glycolytic intermediates (on per carbon basis) stayed mostly steady from 30 min to 60 min (Fig. 1d). This average labeling enrichment normalized to the labeling enrichment of plasma glucose indicates the maximal direct contribution circulating glucose may have to glycolysis^30^. This ratio stayed around 50% (Fig. 1d), showing the direct contribution of plasma glucose was around or slightly less than 50%.

We further assessed the utilization of glucose in primary human neutrophils in an *ex vivo* setting, where freshly isolated peripheral blood neutrophils were assayed in culture media (Fig. 1a). Extracellular glucose labeled slightly less than 50% of the glycolytic intermediates across donors (Fig. 1e), with the dominant label forms reflecting the direct contribution via glycolysis (Supplementary Fig. 1h), and glucose minimally labeled TCA cycle intermediates (Supplementary Fig. 1i). These results show the utilization of glucose is similar *ex vivo* compared to *in vivo* condition, contributing to slightly less than 50% of glycolysis, and there are other important nutrients that contribute significantly to directly supplying the metabolism of primary human neutrophils under resting condition.

To examine these other sources, we next tested the contribution of intracellular glycogen, which recent studies have suggested is important for supplying glycolysis across multiple tissues and for neutrophil survival and functions^5,17,30^. When we inhibited neutrophils’ glycogen utilization by treating them with glycogen phosphorylase inhibitor (GPi), labeling from extracellular glucose became nearly complete (Fig. 1e, Supplementary Fig. 1h). The contribution of glycogen to generating glycolysis intermediates was indicated by the increase in labeling enrichment when glycogenolysis is blocked (+GPi) compared to baseline (-GPi), and we found the contribution of glycogen was ∼50% (Fig. 1f). While glucose and glycogen combined account for most direct nutrient sources, other sources may make a minor contribution (<10%). For instance, we observed significantly less labeled glycerol-3-phosphate (G3P) relative to DHAP both with or without GPi (Supplementary Fig. 1j). The additional unlabeled G3P can be generated from unlabeled sources such as lipid degradation, which can further incorporate into lower glycolysis.

Unlike primary human neutrophils assessed *ex vivo*, which showed a ∼45:50 mixed nutrient preference respectively towards glucose and glycogen, which is consistent with *in vivo* results, we found human neutrophil-like cell line HL-60 cultured *in vitro* primarily depended on extracellular glucose as the direct source for glycolytic intermediates. Extracellular glucose labeled nearly all of the glycolytic intermediates and treatment with GPi had minimal effect (Fig. 1g). This observation is consistent with a recent report in T cells^31^ that found a freshly isolated *ex vivo* model, but not *in vitro* cell models, faithfully reflects *in vivo* metabolic phenotype, likely due to significant metabolic changes that accumulate during prolonged *in vitro* culturing.

### Stimulations induce nutrient preference shift toward primarily using extracellular glucose

As results above revealed that intracellular glycogen, in addition to extracellular glucose, is a major metabolic source for primary human neutrophils at resting state, we next investigated whether such nutrient preference changes when neutrophils are activated by a variety of stimulation signals. Primary human peripheral blood neutrophils were stimulated *ex vivo* with phorbol myristate acetate (PMA), zymosan A (Zym), TNFα, or heat-inactivated *Pseudomonas aeruginosa* (HI PsA; heat inactivation prevents the interference of bacterial metabolism) for 30 min, or LPS for 4 hours, and we traced the incorporation of extracellular U-^13^C-glucose into glycolytic intermediates as an indication of glucose utilization. We observed a significant increase in the labeling of all glycolytic intermediates from less than 50% to 80–90% upon activation with each of these stimuli (Fig. 2a). Similarly, intermediates of the PPP also showed significantly increased labeling from extracellular glucose upon activation (Fig. 2b). These data indicate a substantial nutrient preference shift, with extracellular glucose becoming the dominant direct source for glycolysis and the PPP upon activation.

**Figure 2.**
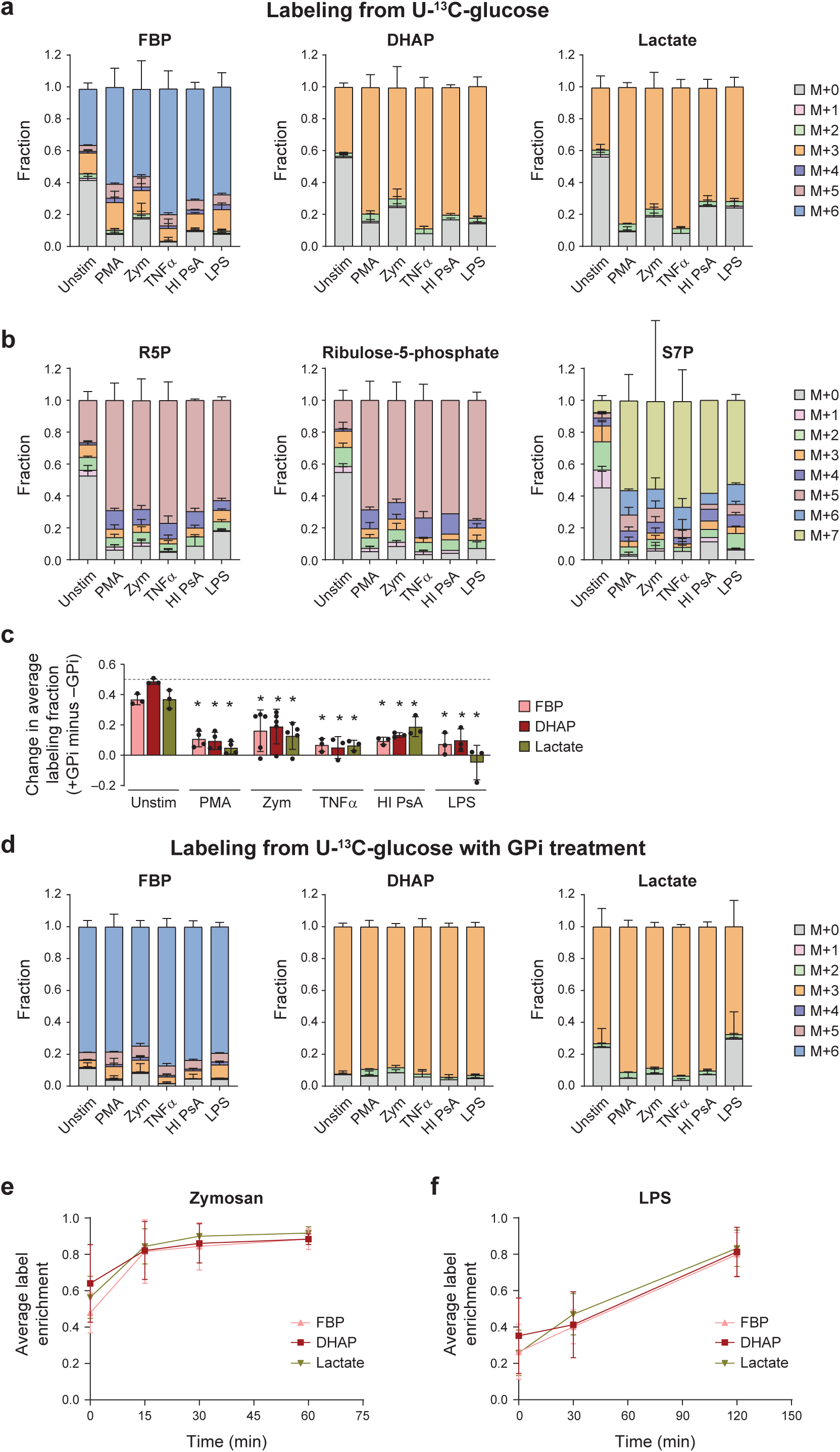
**Nutrient preference shift in neutrophils upon activation.** a, b. Labeling patterns of glycolytic intermediates (a) and pentose phosphate pathway intermediates (b) from U-^13^C-glucose in human peripheral blood neutrophils that are unstimulated (n=5 donors), 30min after stimulation with 100nM PMA (n=4 donors), 300μg/mL zymosan A (n=5 donors), 100ng/mL TNFα (n=3 donors), or heat inactivated *P. aeruginosa* (HI PsA, 10:1 bacteria: neutrophil) (n=3 donors), or 4h after stimulation with 1μg/mL LPS (n= 3 donors). Bars and error bars show mean ± SD. c. Increase in average label enrichment of indicated glycolytic intermediates from U-^13^C-glucose caused by treatment of GPi (20μM), in neutrophils that are unstimulated or stimulated with indicated signals. Bars and error bars show mean ± SD from n=3–5 independent donors, as represented by individual dots. The GPi induced labeling increase of each compound in stimulated neutrophils were compared with that in unstimulated neutrophils by two-tailed, unpaired *t*-test, * indicates p < 0.05. d. Labeling patterns of glycolytic intermediates in neutrophils as described in A, but treated with 20μM GPi. n=3–5 independent donors, as represented by individual dots in panel c. e-f. Dynamic changes of labeling enrichment from U-^13^C-glucose of indicated glycolytic intermediates, following stimulation with 300μg/mL zymosan A (e) or 1μg/mL LPS (f). Bars and error bars show mean ± SD from n=2 (e) or n=3 (f) independent donors.

Mirroring this shift was a substantially reduced contribution of glycogen to glycolysis, as measured by the gain of labeling from glucose when cells were treated with GPi (Fig. 2c). The contribution by other sources beyond glucose and glycogen remained less than 10% in activated neutrophils, as indicated by the fraction of glycolytic intermediates remaining unlabeled from glucose after glycogen degradation was inhibited (Fig. 2d).

While all the investigated stimuli induced a similar nutrient preference shift, the shift occurred with different kinetics. Zymosan induced a very rapid shift, with glucose becoming the dominant source within 15 min after stimulation (Fig. 2e). In contrast, LPS induced such a shift much more slowly, over 2 hours (Fig. 2f).

### The shift to glucose utilization is driven by a substantial increase in glucose uptake by GLUT1

The stimulation-induced nutrient preference shift could be driven by increased glucose utilization or decreased glycogen utilization (Fig. 3a). Therefore, to understand the mechanism for the shift, we first measured glucose consumption rate. A 4- to 6-fold increase in total glucose consumption was observed within 1 hour after stimulation with PMA, zymosan, and TNFα (Fig. 3b). We also found the level of GLUT1 (one of the main glucose transporters in neutrophils^19,32^) on the cell surface increased by ∼5 fold as soon as 30 min after stimulation (Fig. 3c), which would enable the rapid increase in glucose uptake. It has been reported that GLUT1 can be phosphorylated by PKC, and the phosphorylation enhances its localization to the cell surface^33^. Moreover, both zymosan and TNF can activate PKC^34,35^, and PMA is a direct activator of PKC, pointing to a likely mechanism for increased GLUT1 on plasma membrane within 30 min. Indeed, we found that while the total cellular level of GLUT1 remained mostly constant during this time, GLUT1 phosphorylation was increased in these conditions (Fig. 3d).

**Figure 3.**
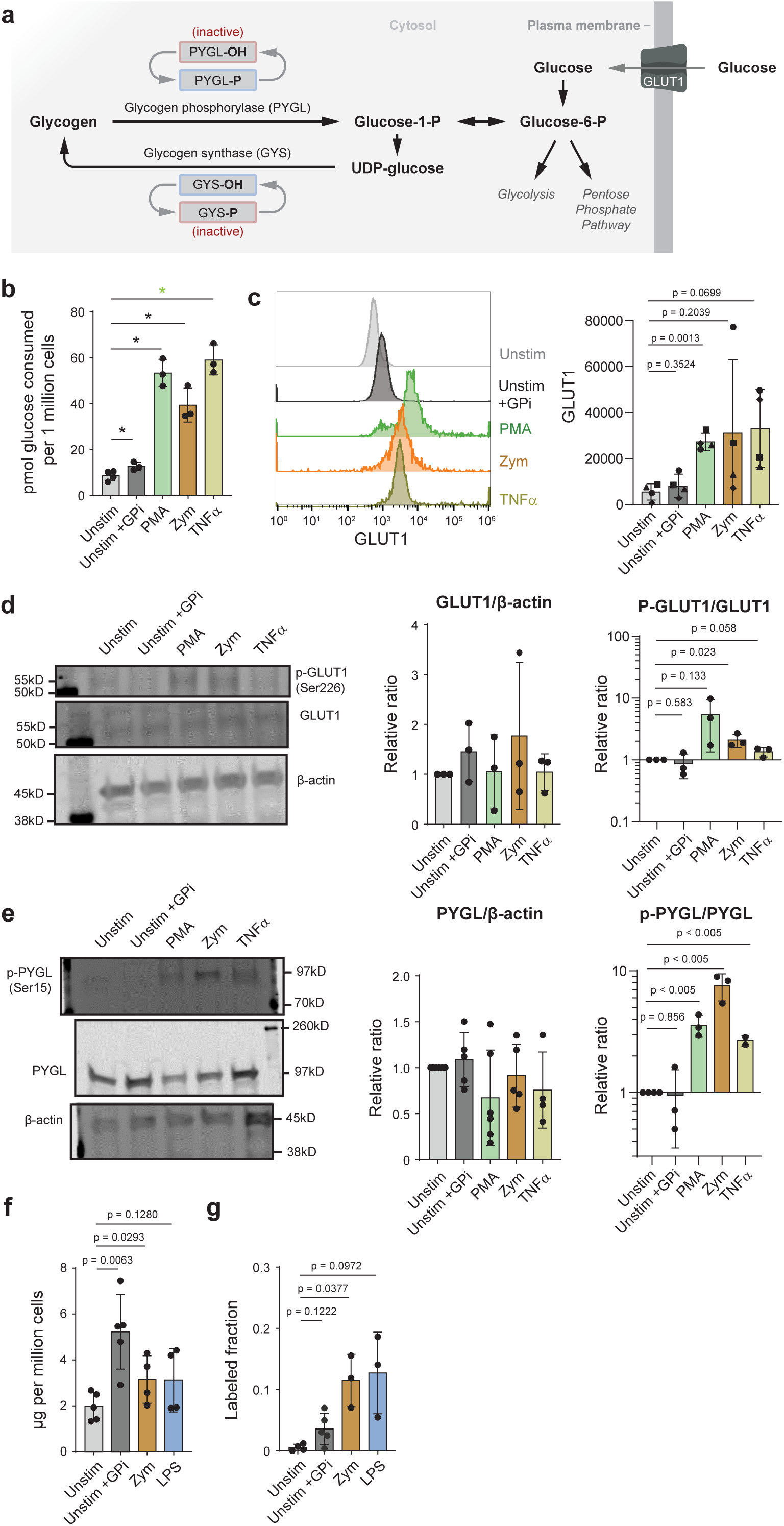
**Changes of glucose uptake and glycogen cycling upon neutrophil activation.** a. Schematic of glucose uptake, glycogen synthesis, and glycogenolysis. Key enzymes are indicated. b. Glucose consumed from the media within 1 hour by unstimulated neutrophils or neutrophils that are treated with indicated stimuli (100nM PMA, 300μg/mL zymosan A, or 100ng/mL TNFα) or inhibitor (20μM GPi). Bars and error bars show mean ± SD from n=3–4 independent donors, as represented by individual dots. Statistical analysis was done by two-tailed, unpaired *t*-test, * indicates p<0.05. c. (Left) Representative flow cytometry result measuring the distribution of GLUT1 level on cell surface within a population of neutrophils that are untreated or treated with indicated stimuli or inhibitor for 30 min. (Right) The experiment was repeated with neutrophils isolated from n=4 independent donors (indicated by different markers). Bars and error bars show mean ± SD of quantified mean GLUT1 level from each donor. Statistical analysis was performed by two-tailed, paired *t*-test. d–e. Immunoblot analysis of (d) total GLUT1 level and phosphorylated GLUT1 (Ser226), and (e) total PYGL and phosphorylated PYGL (Ser15) in neutrophils treated with indicated stimuli or inhibitor for 30min. Representative blots were shown on the left. The experiment was repeated multiple times with neutrophils isolated for different donors (indicated by individual dots), and quantified results were compiled and shown on the right. Bars and error bars show mean ± SD. Statistical analysis performed by two-tailed, unpaired *t*-test. f. Level of glycogen store in human peripheral blood neutrophils that are untreated, treated with GPi (20μM) for 2h, or stimulated with zymosan A (300μg/mL), or LPS (1μg/mL) for 2h. Bars and error bars show mean ± SD from n=4–5 independent donors, as represented by individual dots. p-value determined by two-tailed, paired *t*-test. g. Fraction of labeled glycogen after 2h incubation of neutrophils that are untreated or treated with indicated stimuli or inhibitor in media containing U-^13^C-glucose. Bars and error bars show mean ± SD from n=3–5 independent donors, as represented by individual dots. p-values were determined by two-tailed, paired *t*-test.

Second, as glycogen degradation is regulated largely by the phosphorylation of glycogen phosphorylase (PYGL) (Fig. 3a), we next probed the level and phosphorylation of PYGL. (Phosphorylated PYGL is the active form.) In cells stimulated with PMA, zymosan, and TNFα, there was no significant change in total PYGL level, though phosphorylation of PYGL (Ser15) was significantly increased (Fig. 3e). This data suggests that notwithstanding the relative preference shift toward less glycogen contribution, the gross flux of glycogenolysis is likely still increased upon stimulation -- a more profound increase of glucose uptake can dominate the relative contribution of glycogenolysis.

### Glycogen cycling is increased upon neutrophil activation

Intracellular glycogen is often actively turned over by both degradation and synthesis, a process known as glycogen cycling. Although PYGL is increasingly phosphorylated upon neutrophil activation, indicating increased glycogen degradation activity, the total glycogen storage level in cells trended towards slight increase upon stimulation of zymosan or LPS (Fig. 3f). This observation suggests further increased gross glycogen synthesis flux that results in net balance or accumulation of intracellular glycogen. We measured the labeling of intracellular glycogen after cells are cultured in media containing U-^13^C-glucose for 2 hours, which indicates glycogen turnover during this period. The glycogen labeling increased from <2% in resting state to ∼10% in zymosan-or LPS-stimulated neutrophils within 2 hours (Fig. 3g), and the extent of this increase exceeded the increase of labeling of UDP-glucose (Supplementary Fig. 2a), the precursor of glycogen synthesis. These results suggest that stimulation of neutrophils activate glycogen cycling, including both gross glycogen synthesis and glycogen degradation, and that the net flux favors glycogen synthesis to increase glycogen storage, which may help prepare cells for being recruited to low glucose microenvironments.

### Increased glucose utilization supplies essential metabolic reactions in activated neutrophils

The stimulation of neutrophils triggers a series of immune functions that entail substantial metabolic demands. Greatly increased glucose uptake, along with increased PYGL activity, would increase the production rate of G6P, a common upstream metabolite supplying important biosynthetic and energy producing pathways, including glycolysis, PPP, and TCA cycle. For glycolysis, we found the production of end-product lactate increased substantially over 1 hour after stimulation with PMA, zymosan, or TNFα (Fig. 4a), indicating increased glycolysis rate. Regarding PPP, consistent with our previous observation that PPP flux is greatly upregulated in activated neutrophils to enable effector functions^6^, here we observed PPP intermediates significantly accumulated by 30 min upon stimulation of PMA, zymosan, TNFα, or HI PsA. Only LPS stimulation did not induce such change by 4 hours (Fig. 4b). To understand how changes in nutrient utilization support such upregulation of PPP upon activation, we limited the utilization of glucose or glycogen. Withdrawing extracellular glucose significantly reduced the accumulation of pentose phosphate upon stimulation. In contrast, inhibiting glycogen degradation only had a minor impact on the accumulation of PPP intermediates, and the combination of glucose withdrawal and glycogen utilization inhibition had the strongest effect (Fig. 4c). Consistent with glucose labeling showing that extracellular glucose became the dominant source of PPP upon activation (Fig. 2b), this result suggests increased glucose utilization, but not glycogen utilization, is essential to support the upregulation of PPP.

**Figure 4.**
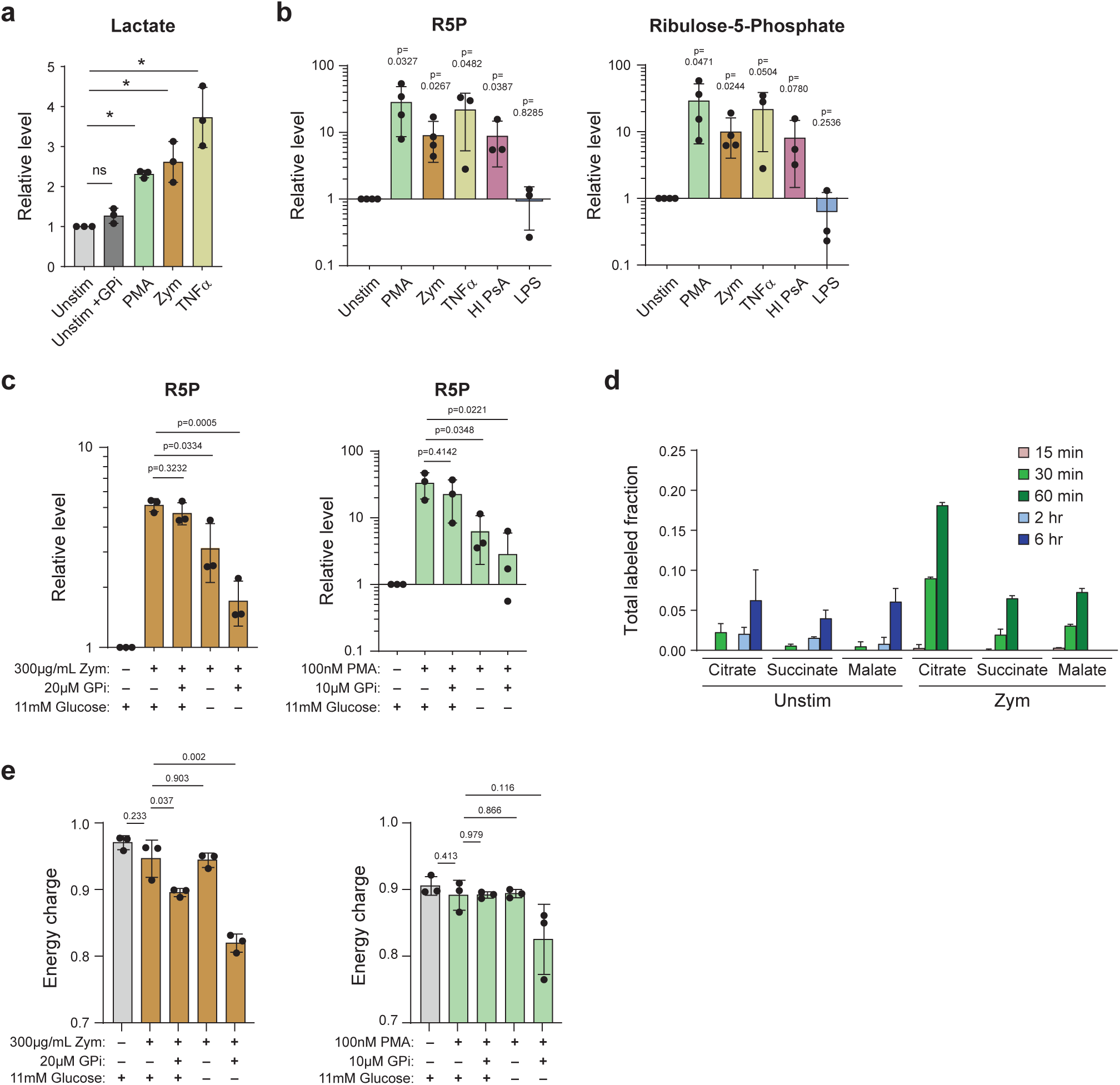
**Increased nutrient utilization upon activation supplies various metabolic pathways.** a. Lactate accumulation in spent media of neutrophils that are untreated or treated with GPi or indicated stimuli after 1 hour. Bars and error bars show mean ± SD from n=3 independent donors, as represented by individual dots. * indicates p < 0.05 by two-tailed, unpaired *t*-test. b. Relative ribose-5-phosphate (R5P) and ribulose-5-phosphate abundance in neutrophils that are unstimulated or stimulated with 100nM PMA, 300μg/mL zymosan A, 100ng/mL TNFα, or heat inactivated *P. aeruginosa* (HI PsA) for 30 min, or with 1μg/mL LPs for 4 hours. Bars and error bars show mean ± SD from n=3 independent donors, as represented by individual dots. p-values were determined by two-tailed, unpaired *t*-test comparing stimulated condition to unstimulated. c. Relative R5P abundance (normalized to unstimulated condition) in neutrophils stimulated with zymosan A or PMA for 30min, cultured in media with or without glucose, with or without the treatment of GPi. Bars and error bars show mean ± SD from n=3 independent donors, as represented by individual dots. p-values were determined by two-tailed, unpaired *t*-test comparing each nutrient perturbed condition (with glucose withdrawal or with GPi treatment) to stimulated neutrophils cultured in standard culture condition (with glucose, with no GPi). d. Total labeled fraction of TCA cycle metabolites in neutrophils (adhered to tissue culture plate) that are unstimulated or stimulated with 300μg/mL zymosan A after incubation with media containing U-^13^C-glucose for indicated time. e. Energy charge in unstimulated neutrophils, or neutrophils stimulated with zymosan A or PMA for 30 min, cultured in media with indicated nutrient perturbation (with or without glucose, with or without GPi treatment). Bars and error bars show mean ± SD from n=3 independent donors, as represented by individual dots. p-values were determined by two-tailed, unpaired *t*-test.

Next, we assessed how the TCA cycle is altered by increased glucose utilization upon neutrophil activation. In neutrophils under resting condition, labeled glucose was incorporated into the TCA cycle slowly and to a minor level, with most TCA cycle intermediates only labeled ∼5% by 6 hours (Fig. 4d). The low baseline activity of TCA cycle is consistent with previous literature suggesting that neutrophils have few classic mitochondria^4,36^. Upon activation with zymosan, we found a rapid and substantial increase in TCA cycle labeling from glucose, with citrate labeling up to 20% within 1 hour (Fig. 4d), which exceeds the extent of increased glycolytic intermediates labeling. This result suggests that upon activation, increased flux was sent through TCA cycle, supported by increased glucose utilization. Nonetheless, the overall fraction of TCA cycle labeling was relatively small (<20%) within the short period after stimulation, suggesting TCA cycle turnover is a relatively slow process for glucose utilization, and there may be other important sources beyond glycolytic products feeding into TCA cycle in neutrophils. Consistent with this, neither glucose withdrawal nor glycogenolysis inhibition within a short period significantly reduced the level of TCA cycle intermediates (Supplementary Fig. 2b).

Despite the substantial metabolic demands triggered by neutrophil activation, cellular energy status was generally well-maintained upon stimulation (Fig. 4e), suggesting the increased glucose utilization, as well as gross glycogen utilization, is sufficient to support the increased bioenergetic demand upon activation. In testing the dependence of cellular energetics on glucose or glycogen utilization, we found that inhibiting glycogen utilization or withdrawing extracellular glucose alone did not substantially reduce energy charge in stimulated neutrophils, while inhibiting both noticeably reduced energy charge (Fig. 4e, Supplementary Fig. 2c). This finding suggests that with increased capacity to utilize both glucose and glycogen, neutrophils can flexibly use either source to maintain energy status, which can in turn provide the basis for their metabolic flexibility to support cell functions under different nutrient conditions.

### Utilization of extracellular glucose is required for oxidative burst

Results above revealed the shift in nutrient utilization and the downstream changes in metabolic pathways induced by neutrophil stimulation. The reprogramming of cellular metabolism plays a key role in supporting immune functions neutrophils perform quickly upon activation. We next investigated how suppressing or priming such activation-induced shift towards glucose utilization impacts neutrophil functions, and examined the extent to which various neutrophil functions depend on the two major metabolic sources identified, glucose and glycogen. To do so, we measured the changes in neutrophil functions after acutely limiting the utilization of either one or both nutrient sources.

As an important control, we first examined how such nutrient perturbations impact cell viability, and found that blocking glycogenolysis with glycogen phosphorylation inhibitor, glucose withdrawal, or the combination of both, did not reduce cell viability within a period of a few hours (Supplementary Fig. 3a). A day after primary neutrophil isolation, a large portion of the cells died across all of these nutrient-limited conditions, similar to the control condition (Supplementary Fig. 3b). This is consistent with neutrophils’ short-lived nature. For assaying cell functions, we focused on the first few hours after neutrophil isolation as neutrophils are the first responders in innate immunity and perform their functions rapidly. During this period, cells remained equally highly viable across all nutrient conditions (Supplementary Fig. 3a).

Oxidative burst is the production of reactive oxygen species for pathogen killing, which can be induced by PMA or zymosan stimulation. Inhibiting glycogen utilization had minimal impact on oxidative burst induced by PMA or zymosan, while inhibiting extracellular glucose utilization by inhibiting hexokinase (using 2DG) or glucose withdrawal greatly suppressed the oxidative burst (Fig. 5a). Limiting glucose utilization not only greatly reduced the overall intensity of burst (Fig. 5b), but also quickened the decline of the oxidative burst, particularly for lower dose stimulation, which induces a slower burst (Fig. 5c). This finding is consistent with neutrophils relying heavily on glycogen utilization at baseline but switching to primarily using glucose soon after stimulation. Thus, when extracellular glucose is absent, oxidative burst is still able to start but soon becomes limited. The effect of inhibiting glucose and glycogen utilization appears to be additive; in the condition with no glucose and with GPi treatment, cells have the lowest amount of burst (Fig. 5 a,b). Together, these results showed the oxidative burst in activated neutrophils primarily depends on high glucose utilization, with glycogen providing a very minor contribution but unable to replace the role of extracellular glucose. The shift towards glucose utilization is required for oxidative burst.

**Figure 5.**
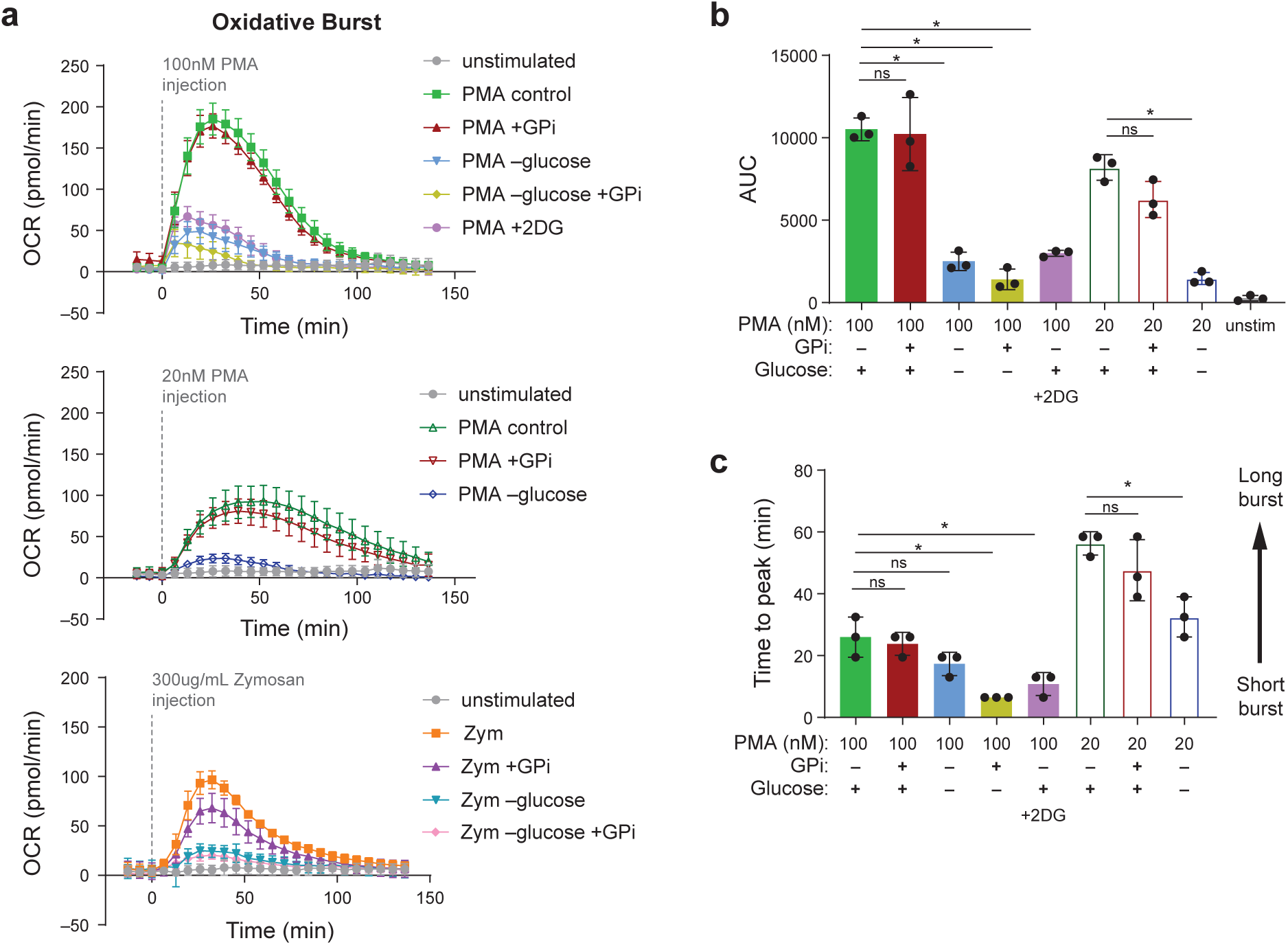
**Nutrient dependence of oxidative burst.** a. Oxidative burst induced by 100nM or 20nM PMA or 300ug/mL zymosan A in indicated nutrient conditions. “Control” – standard culture media; “-glucose” – media with no glucose; “+ GPi”, treated with 20μM GPi; “+2DG” – treated with 1mM 2DG; “-glucose+GPi” – cultured in media with no glucose and treated with 20μM GPi; “unstimulated”—neutrophils cultured in standard media without stimuli. The oxidative burst curves shown are a representative result from one neutrophil donor, dot and error bar show mean ± SD from technical replicates (n=8). b–c. The experiment described in (a) was repeated with peripheral blood neutrophils isolated from n=3 independent donors, the area under curve (AUC) over the oxidative burst was quantified from results from each donor and shown in (b), and the time from stimulation to the peak of oxidative burst was quantified and shown in (c). Bars and error bars show mean ± SD. Dots represent result from individual donors. * indicates p < 0.05, ns indicate non-significant (p>0.05) by two-tailed, unpaired *t*-test.

### Neutrophil extracellular trap release and phagocytosis can be flexibly supported by either glucose or glycogen

NET is a network of decondensed DNA mixed with granule enzymes that neutrophils release upon activation for trapping and killing pathogens. Contrasting with oxidative burst, inhibiting the utilization of either extracellular glucose or intracellular glycogen only slightly decreased NET release, as measured by the amount of extracellular DNA (Fig. 6a, b). However, inhibiting both completely abolished NET release (Fig. 6b). This finding suggests neutrophils have great flexibility to use either nutrient source to support NETosis, but at least one of these sources is required and cannot be compensated by other alternative sources.

**Figure 6.**
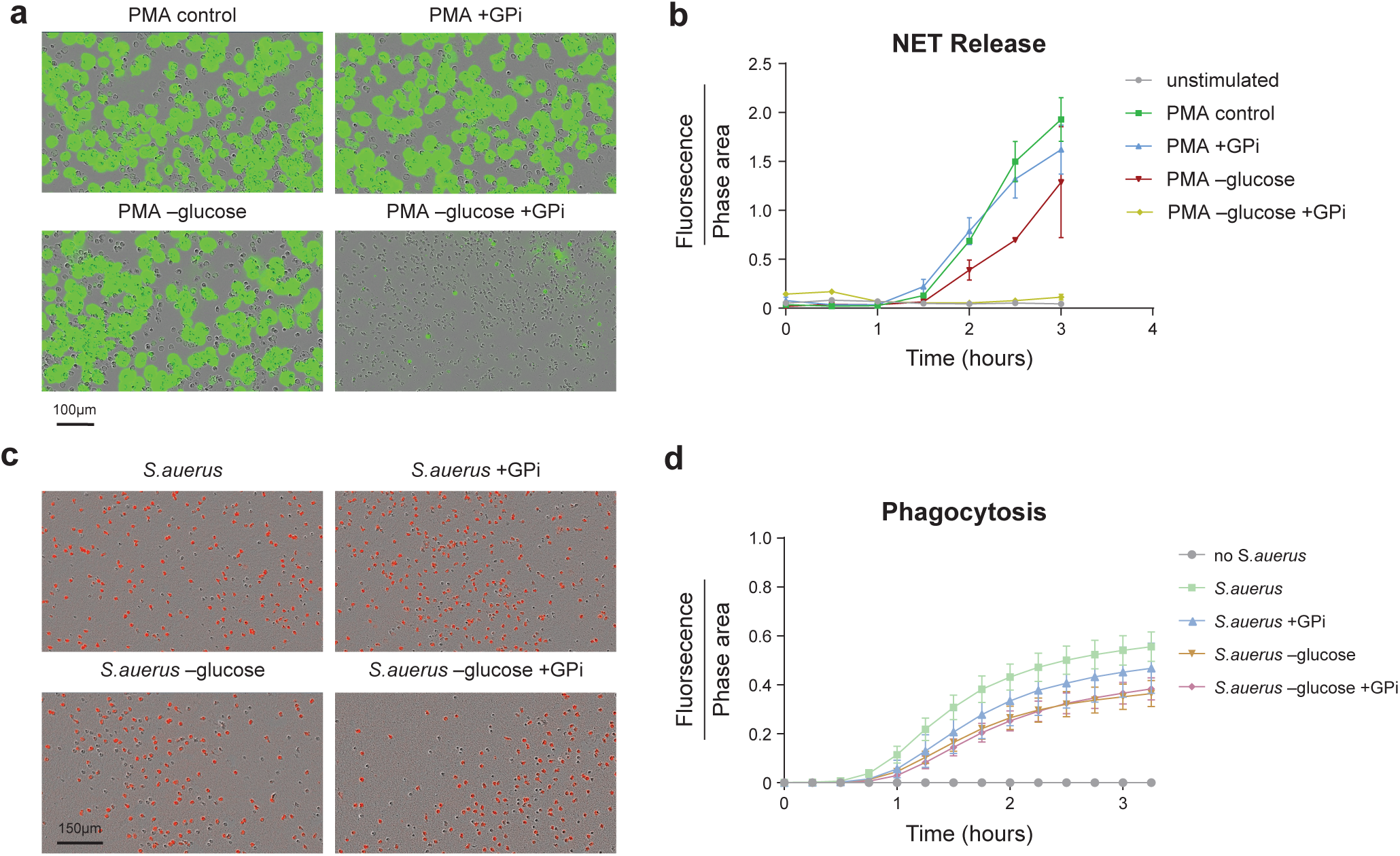
**Nutrient dependence of neutrophil extracellular trap release and phagocytosis.** a. Representative images of NETs released by peripheral blood neutrophils cultured in indicated nutrient conditions 2.5 hour after stimulation with 100nM PMA. Green fluorescent signal from extracellular DNA (visualized by Cytotox Green dye) indicates NETs. b. NET release, quantified from images represented in (a), over a time course after stimulation with PMA in peripheral blood neutrophils cultured in indicated nutrient conditions. Dot and error bar show mean ± SD from n=3 independent donors. c. Representative images of phagocytosis by peripheral blood neutrophils cultured in indicated nutrient conditions 2.5 hour after incubation with pHrodo (red fluorescent)- labeled *S. aureus* bioparticles. d. Phagocytosis of *S. aureus* over a time course, quantified from images represented in (c), by peripheral blood neutrophils cultured in indicated nutrient conditions. Curves show representative result. Dot and error bar indicate mean ± SD from n=9 technical replicates. Trends have been confirmed by independent experiments using neutrophils isolated from three donors.

Similar to NET release, phagocytosis of bacteria also showed flexibility in nutrient dependence. Cells could still largely perform phagocytosis when either glucose or glycogen utilization, or both, was limited, although the phagocytosis rate was reduced when either nutrient supply was limited (Fig. 6c, d). For both NET release and phagocytosis, the dependence on extracellular glucose is slightly higher than glycogen.

### Shifting away from glycogen utilization increases neutrophil migration and fungal control

Upon sensing signals associated with infection or inflammation, neutrophils can quickly migrate towards the site. We next examined how the shift from glycogen to glucose utilization impacts neutrophil migration. Migration towards chemoattractant C5a was greatly inhibited by the withdrawal of extracellular glucose, suggesting high dependence of migration on glucose utilization. Conversely, inhibiting glycogen degradation with GPi, which would force cells shift to primarily rely on extracellular glucose (Fig. 1e), significantly increased neutrophil migration (Fig. 7a). Interestingly, when glycogen degradation inhibition is combined with glucose withdrawal, we observed an even greater suppression of neutrophil migration than glucose withdrawal alone (Fig. 7a). It is possible that the metabolic shift towards glucose utilization can promote migration; however, cells’ ability to migrate also depends on sufficient cellular energy charge, which becomes limited when inhibiting glycogen utilization in an environment lacking extracellular glucose.

**Figure 7.**
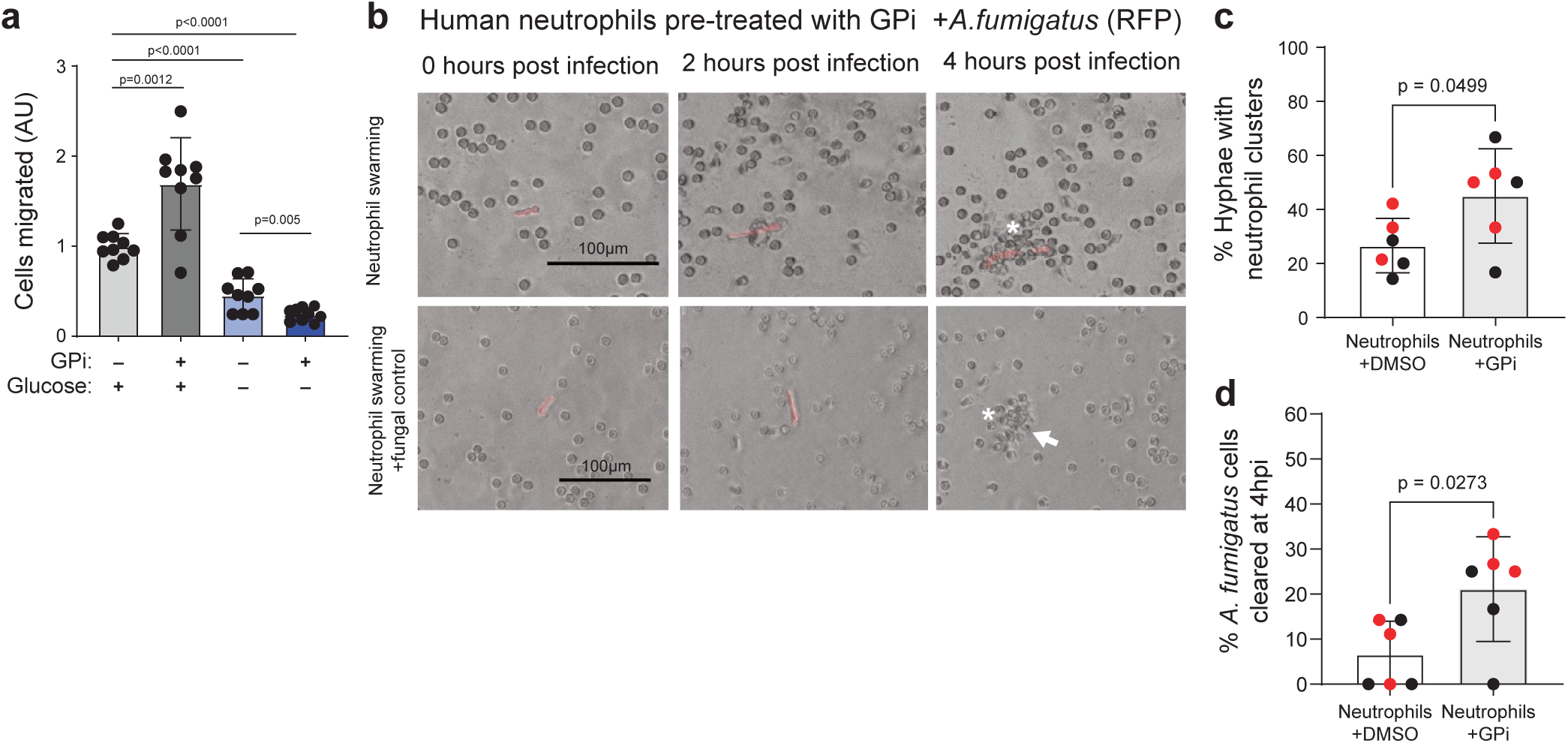
**Nutrient dependence of neutrophil migration and pathogen killing.** a. Migration of neutrophils through transwell towards 2nM C5a in indicated nutrient condition (with or without glucose, with or without GPi treatment). Bars and error bars show mean ± SD of pooled technical replicates (represented by dots) from three individual donors. p-values were determined by two-tailed, unpaired *t*-test. b. Representative images of human peripheral blood neutrophils (pre-treated with 20μM GPi for 1h before co-incubation with *A. fumigatus* in standard culture media) effectively clustering around and killing *A. fumigatus* hyphae. (Red fluorescent signal is from RFP expressing *A. fumigatus.* White asterisks indicate neutrophil cluster formation. White arrow indicates fungal cells that have been killed, as indicated by loss of cytosolic RFP signal.) c. Percent of *A. fumigatus* hyphae with neutrophil clusters (5 or more neutrophils) around them after 4 hour co-incubation with untreated primary human neutrophils or neutrophils pretreated with 20uM GPi. d. Percent of *A. fumigatus* cells cleared after 4 hour co-incubation with untreated primary human neutrophils or neutrophils pretreated with 20μM GPi. c–d. Bars and error bars show mean ± SD of pooled technical replicates (represented by individual dots) from two individual donors (indicated by different colors). Statistical analysis was performed by two-tailed, unpaired *t*-test. (n=70 total fungal cells analyzed for the DMSO treated condition and n=45 total fungal cells analyzed for the GPi-treated condition).

Given the intriguing observation that inhibition of glycogenolysis promotes neutrophil migration towards chemoattractant, we further assessed the impact of nutrient utilization shift on neutrophil migration and its downstream impact in pathogen control in the setting of fungal infection with *Aspergillus fumigatus*^37^-- Neutrophils play a significant role in controlling the growth of the invasive pathogen *A. fumigatus,* and coordinated swarming towards the pathogen enables effective fungal control. In this experiment, to avoid the direct impact of GPi treatment on *A. fumigatus*, we pre-treated primary human neutrophils with GPi for 1 hour, then removed the GPi and co-incubated neutrophils with fungal germlings in fresh culture media. The inhibitory effect of neutrophil glycogenolysis by GPi is sustained up to 4 hours after pre-treatment (Supplementary Fig. 4a). Consistently, the effect of GPi treatment on inhibiting neutrophil migration towards C5a can also be sustained after an acute pre-treatment (Supplementary Fig. 4b). When co-cultured with *A. fumigatus*, neutrophils swarmed towards and clustered around fungal pathogens (Fig. 7b). The percentage of neutrophil clusters around fungal hyphae was significantly increased when neutrophils were pre-treated with GPi, suggesting increased neutrophil migration towards fungal pathogens upon glycogenolysis inhibition (Fig. 7c). Consistent the increased neutrophil clustering around developing *A. fumigatus* hyphae and germlings (Fig. 7b,c), the fraction of *A. fumigatus* cells that were cleared after 4 hours co-incubation with neutrophils was significantly increased when the neutrophils were pre-treated with GPi (Fig. 7d). This result further shows that shifting neutrophil away from glycogen utilization promotes fungal control.

## Discussion

This work quantitatively evaluated the contributions of various nutrients in neutrophil metabolism under resting and activated states. Glycogen was surprisingly found to be a major metabolic source, with contribution equal to or greater than glucose, particularly at resting state. Neutrophils can actively turn over glycogen (a turnover that we found is upregulated upon cell activation). The synthesis of glycogen uses UDP-glucose as the direct substrate, which is synthesized from G6P. Recent work which found that neutrophils express key gluconeogenic enzymes^5^ has raised the potential role of gluconeogenesis to supply glycogen storage. We found that at gross flux level, a small fraction of glycolytic intermediates does not directly arise from glucose or glycogen, which may result from gross flux in the direction of gluconeogenesis. However, such contribution is minor. At the net flux level, supplying glycogen with gluconeogenesis requires ATP production from non-glycolytic pathways to support bioenergetics. Our observation is that, consistent with previous reports, TCA cycle activity in resting neutrophils is very low to support energy production. Therefore, overall, glucose is the ultimate source for neutrophil metabolism and energy supply.

However, at gross flux level, a large fraction of the glucose is converted to glycogen first, and glycogen acts as a major direct source for neutrophil metabolism at resting state.

The question then becomes why primary neutrophils actively cycle between glucose and glycogen, rather than directly use extracellular glucose like some other cells, or even the neutrophil-like cell line. Cycling glucose with glycogen storage can provide a buffer of fluctuating glucose level. Primary neutrophils need to be actively recruited from circulation to the site of infection, where they perform important functions. Such sites often have low microenvironmental glucose^38^, and therefore high capacity to use glycogen storage can provide neutrophils extra metabolic flexibility, even though extracellular glucose is the preferred source for activated neutrophil when available. On the other hand, relying on glycogen cycling rather than just using extracellular glucose directly may also better buffer neutrophils from potential dysregulating effects of high extracellular glucose. For example, hyperglycemia is detrimental during sepsis, and hyperglycemia in diabetes appears to be closely associated with multiple hyper-active phenotypes in neutrophils and altered wound healing^39–41^. These effects of high glucose mirror our observation that shifting towards glucose utilization is associated with activation of certain neutrophil functions. The idea that the ability to cycle between glucose and glycogen is critical for neutrophils is further supported by clinical evidence in glycogen storage disease (GSD). Low neutrophil numbers and functional defects in neutrophils are among the most important symptoms of some subtypes of GSD, particularly GSD-Ib^42–45^. Interestingly, some other forms, such as GSD-Ia, have no clinical reports of similar neutrophil dysfunction^46^. GSD-1b is characterized by a deficiency in G6P transport through the endoplasmic reticulum membrane. This further suggests that not only is general glycogen cycling important for neutrophils, but there may also be subcellular compartment-specific metabolism that allow neutrophils to channel glucose or glycogen utilization differently, which is critical to neutrophil function. The compartment-specific role may also contribute to some of the function-specific dependence on glucose or glycogen we report here, and requires further investigation.

While in general active cycling between glucose and glycogen can provide more buffering and metabolic flexibility, our results here also reveal that activation prompts neutrophils to utilizing extracellular glucose primarily, and that diverting the nutrient utilization can significantly alter their functions. In general, suppressing the activation-induced shift towards glucose utilization more significantly suppressed functions performed by activated neutrophils. Intriguingly, priming the nutrient preference shift by treatment with glycogenolysis inhibitor, which we found increased glucose utilization both in relative contribution and in absolute rate, can promote some important functions, including neutrophil migration and fungal control. Indeed, we found the exact effect of modulating nutrient metabolism on neutrophil functions is function specific, as these functions vary in their flexibility to use glucose or glycogen to compensate for each other. Notably, the oxidative burst depends on the shift to extracellular glucose utilization indispensably, whereas NET release can be very flexibly supported by either glucose or glycogen.

This function-specific nutrient dependence likely stems from the specific type, size, and kinetic requirement of any given function’s major metabolic requirements: in short, different functions entail different metabolic demands. Oxidative burst, for instance, requires neutrophils to generate a large amount of NADPH in a short period of time to fuel the production of reactive oxygen species via NADPH oxidase. We have previously shown that this rapid NADPH production depends on substantial upregulation of the PPP^6^. Here we found the PPP upregulation is primarily supported by glucose, which cannot be replaced by glycogen (Fig. 4c). This is likely because activated neutrophils can use extracellular glucose to supply substrates at much higher gross flux than they can utilize glycogen (as indicated by the labeling results), even though the ability to use both nutrient sources is activated. Therefore, glucose, the fast-accessible source, has an irreplaceable role in supporting the high metabolic demands of oxidative burst, and the effects of glucose withdrawal or glycogen degradation inhibition on oxidative burst mirror their effects on PPP (Fig. 5). By contrast, other functions may be associated with metabolic demands that are not as kinetically limited (e.g., NET release only needs a low level of reactive oxygen species to signal for its activation^47^), or they may primarily require energy supply. We found that activated neutrophils can largely maintain their energy status with either glucose or glycogen, as long as at least one of the sources is available (Fig. 4e). Such functions with moderate demands are more likely to have high flexibility to be supported by either nutrient, and such metabolic flexibility is further ensured by neutrophils’ increased ability to use both glucose and glycogen upon activation thanks to the rapid phosphorylation and activation of GLUT1 and PYGL.

These findings of neutrophils’ nutrient metabolism, and its significant and specific impacts on immune functions, have important implications for health, given the crucial role of proper neutrophil functions in a broad range of contexts. While promoting pathogen-eliminating functions is essential for combating against infection, over-activation of functions such as oxidative burst can cause tissue damage, and lowering pro-inflammation response can alleviate inflammatory conditions. Therefore, understanding neutrophils’ specific metabolism–function connections is meaningful both for discerning how altered nutrient metabolism may contribute to diseases and for targeting nutrient metabolism to modulate specific functions for treatment. Indeed, leveraging specific metabolic manipulation to shape immune response has shown promise in other immune cells. For instance, elevated glucose uptake has been recognized a hallmark of activated T cells, B cells, and macrophages, and therefore has been probed as a therapeutic target for various diseases^48,49^. However, equivalent research in neutrophils had been significantly lagging. Providing an important foundation, this study reveals the quantitative contribution of various nutrients in the metabolism of human neutrophils, revealed the reprograming of nutrient metabolism upon neutrophil activation, and elucidates how altering nutrient utilization impacts specific neutrophil functions. Future research is needed to further understand questions in two areas: first, what exact molecular mechanisms underlie the specific connections between nutrient metabolism and neutrophil functions (e.g., what mechanisms explain why steering neutrophils away from using glycogen promotes migration), and second, whether alteration in nutrient levels in specific biological environments impacts progression of disease through changing neutrophil metabolism.

## Methods

### *In vivo* isotopic tracing

Healthy adults with no previous history of diabetes received infusions of U-^13^C-glucose after informed consent was obtained following the protocol approved by the University of Wisconsin Institutional Review Board (2020-1008-CP001). Sterile and endotoxin-free U-^13^C-glucose was administered as a bolus of 8 grams over 10 min followed by an 8 g/hour infusion over 2 hours. Peripheral blood samples were collected at various time points into EDTA coated tubes, and metabolites were extracted from plasma and neutrophils. Plasma was collected by spinning at 1000 x g for 10 min., and plasma metabolites was extracted by adding 50 ul plasma into 200 ul of 40:40:20 acetonitrile/methanol/water (40:40:20 v:v:v). Neutrophils were isolated from peripheral blood collected prior to infusion and at 30 and 60 min following the start of infusion (method see “isolation of human peripheral blood neutrophils” below), then pelleted by spinning at 500*g* for 2 min. Metabolites were extracted from neutrophil pellets by addition of LC-MS grade 40:40:40 acetonitrile/methanol/water (40:40:20 v:v:v). Samples were spun at 20,627*g* for 5 min at 4 °C to remove any insoluble debris prior to analysis on liquid chromatography-mass spectrometry (LC-MS) (see below for details).

To measure plasma glycerol, which does not ionize well without derivatization, it was first enzymatically converted into glycerol-3-phosphate prior to analysis on LC-MS^50^. Briefly, plasma samples were incubated with glucokinase (Sigma-Aldrich G6278) for 10 min at room temperature in buffer (5mM Tris-HCl, 50mM NaCl, 10mM MgCl2, 5mM ATP). After incubation, reaction was quenched with LC-MS grade 40:40:40 acetonitrile/methanol/water +0.1% formic acid (40:40:20 v:v:v) and neutralized with 15% ammonium bicarbonate. Samples were spun at 16,000 x g for 10 mi at 4 °C prior to analysis on LC-MS.

### Isolation, culture and stimulation of human peripheral blood neutrophils

Human neutrophils were isolated from peripheral blood freshly collected from healthy donors, following the protocol approved by the University of Wisconsin Institutional Review Board (protocol no. 2019-1031-CP001). Informed consent was obtained from all participants.

Neutrophils were isolated using the MACSxpress Whole Blood Neutrophil Isolation Kit (Miltenyi Biotec, no. 130-104-434) followed by erythrocyte depletion (Miltenyi Biotec, no. 130-098-196) according to the manufacturer’s instructions. Purity of isolated cells was verified by flow cytometry as previously established^6^.

For *ex vivo* experiments, neutrophils were spun down at 300*g* for 5 min, and resuspended in RPMI 1640 (VWR, no. VWRL0106-0500) supplemented with 5 mM L-glutamine (Fisher Scientific, no. BP379-100) and 0.1% Human Serum Albumin (Lee Biosolutions, no. 101-15) and kept at 37 °C in incubators under 5% CO2. For experiments with glucose withdrawal, the neutrophils were cultured in glucose depleted media: RPMI 1640 (VWR, no. IC091646854) supplemented with 5 mM L-glutamine (Fisher Scientific, no. BP379-100) and 0.1% Human Serum Albumin (Lee Biosolutions, no. 101-15).

To activate neutrophils, between 1.5 and 3 million neutrophils were aliquoted into 1.5-ml Eppendorf tubes. Cells were stimulated with the following stimuli as indicated in each experiment. PMA (Cayman Chemical, no. 10008014) was added to cell suspensions at the indicated concentrations. *P. aeruginosa* (PAK strain) were cultured overnight, sub-cultured until mid-exponential phase then opsonized in human serum (Sigma-Aldrich, no. H4522) for 30 min. Opsonized PsA was spun down at 10,000*g* for 1 min, washed and resuspended in PBS, then heat inactivated at 95 °C for 5 min before addition to neutrophil suspension at 10:1 bacterial/neutrophil cells. Zymosan A was opsonized by incubation with human serum (Sigma-Aldrich, no. H4522) at 37 °C for 30 min, washed twice and resuspended in PBS then added to neutrophils at the indicated concentration for stimulation. TNF-α (R&D systems, no. 210-TA-020) was dissolved in PBS to make a stock solution and added to neutrophils for stimulation. LPS (E. coli O111:B4, Sigma) was dissolved in PBS to make a stock solution then sonicated in a water bath for 10 min prior to addition to cell suspensions. In experiments involving glycogen phosphorylase inhibitor (GPi), 2-chloro-4,5-difluoro-N-[[[2-methoxy-5-[[(methylamino)carbonyl]amino]phenyl]amino] carbonyl]-benzamide (Cayman Chemical, no. 17578) were used, and GPi was added concurrently with stimulation unless otherwise noted at the indicated concentrations.

### Culture of HL-60 cells

HL-60 cells were cultured in RPMI with 5 mM glutamine, 25 mM HEPES, 15% fetal bovine serum (FBS) and 1% penicillin/streptomycin. To differentiate HL-60 to a neutrophil-like state, cells were transferred to differentiation medium (RPMI with 1.3% DMSO, 5 mM glutamine, 25 mM HEPES, 9% FBS and 1% penicillin/streptomycin) for 6 days, with medium change on day 3. After 6 days, differentiated neutrophil-like cells were verified by flow cytometry as previously established^6^ and were used in experiments in culture media without DMSO.

### *Ex vivo* and *in vitro* isotopic tracing

Cells were cultured in labeled medium, which is chemically identical to regular culture medium, but with stable isotope tracer U-^13^C-glucose (Cambridge Isotope, no. CLM-1396) substituting regular glucose at the same concentration, for indicated time. For labeling experiments of stimulated neutrophils, the cells are switched to labeled medium at the same time stimuli (PMA, Zymosan A, TNF, or HI PsA) was added. The exception was LPS stimulation, neutrophils were stimulated with 1ug/mL LPS for 3 hours in unlabeled glucose media prior to switching to U-^13^C-gluose media for an additional hour for a total of 4 hours of stimulation, but only 1 hour labeling, to avoid significant labeling of glycogen due to prolonged culturing in U-^13^C-gluose which complicate result interpretation.

### Metabolomics

To extract intracellular metabolites, culture medium was removed and neutrophils were immediately washed with PBS. Each 2 million pelleted neutrophils were extracted with 150 µl of cold LC–MS-grade acetonitrile/methanol/water (40:40:20 v:v:v), and samples were spun at 20,627*g* for 5 min at 4 C to remove any insoluble debris. To analyze extracellular metabolites, spent media samples were extracted with 4× volume of LC–MS grade methanol, and protein pellets were removed by centrifugation. Supernatant containing small molecules was further diluted with either LC–MS grade H_2_O or acetonitrile/methanol/water (40:40:20 v:v:v) for different LC–MS analysis methods. Soluble metabolite samples were analyzed with a Thermo Q-Exactive mass spectrometer coupled to a Vanquish Horizon Ultra-High Performance Liquid Chromatograph, using the following two analytical methods. (1) Samples in extraction solvent were directly loaded on to LC–MS, then separated on a 2.1 × 150mm Xbridge BEH Amide (2.5 μm) Column (Waters) using a gradient of solvent A (95% H_2_O, 5% ACN, 20 mM NH_4_AC, 20 mM NH_4_OH) and solvent B (20% H_2_O, 80% ACN, 20 mM NH_4_AC, 20 mM NH_4_OH). The gradient used was 0 min, 100% B; 3 min, 100% B; 3.2 min, 90% B; 6.2 min, 90% B; 6.5 min, 80% B; 10.5 min, 80% B; 10.7 min, 70% B; 13.5 min, 70% B; 13.7 min, 45% B; 16 min, 45% B; 16.5 min, 100% B; 22 min, 100% B. The flow rate was 0.3 ml min^−1^ and column temperature 30 °C. Analytes were measure by MS using full scan. (2) Samples were dried under N_2_ flow and resuspended in LC–MS-grade water as loading solvent. Metabolites were separated on a 2.1 × 100mm, 1.7 µM Acquity UPLC BEH C18 Column (Waters) with a gradient of solvent A (97:3 H_2_O/methanol, 10 mM TBA, 9 mM acetate, pH 8.2) and solvent B (100% methanol). The gradient was: 0 min, 5% B; 2.5 min, 5% B; 17 min, 95% B; 21 min, 95% B; 21.5 min, 5% B. Flow rate was 0.2 ml min^−1^. Data were collected with full scan. Identification of metabolites reported here was based on exact *m/z* and retention time that were determined with chemical standards. Data were collected with Xcalibur 4.0 software and analyzed with Maven. For all labeling experiments, natural ^13^C abundance was corrected from raw data. Due to a limited cell number from each neutrophil donor not all the simulation conditions could be done on the same donor, so for determining metabolite abundance, LC-MS data was normalized to resting condition per experiment, and the relative changes induced by stimulation was reported across different donors.

Average labeling enrichment for selected metabolites on a per-carbon basis is calculated based on *Equation 1*.

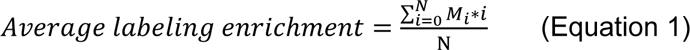

N is the total number of carbon in the compound. M_i_ is the fraction of the compound with i-labeled carbon.

To determine energy charge, relative metabolite abundance of AMP, ADP, and ATP was quantified from ion count in LC-MS analysis based on external calibration curves obtained by measuring a series of AMP, ADP, and ATP standards at various concentration using the same LC-MS method. Energy charge is calculated based on *Equation 2*.

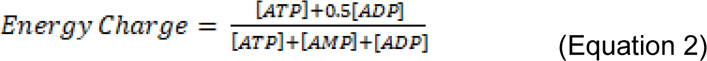

To quantify the level of glucose in plasma, isotopically labeled internal standard was added to plasma samples, and glucose level was qualified based on the ratio between signals from sample and from the standard of known concentration.

### Flow cytometry

To measure levels of neutrophil surface marker and GLUT1, 5×10^5^ neutrophils were aliquoted. Following 15 minutes of Fc-block, primary antibodies, and live/dead dye, Ghost Dye Violet 450 (Tonbo, no.13-0863), were added and incubated at room temperature for 30 min. Primary antibodies: CD11b (PE) (Biolegend, no. 301305), GLUT-1 (FITC) (Alomone Labs, AGT-041-F). After staining, cells were pelleted at 500*g* for 2 minutes and fixed in 0.4% paraformaldehyde at room temperature for 10 min. Cells were then pelleted at 500*g* for 2 min and resuspended in PBS for flow cytometric analysis. Samples were acquired on an Attune NxT V6 (ThermoFisher). Analysis was completed using FlowJo software. Cells were gated for viability and CD11B expression prior to GLUT1 expression gating. A representative gating method on unactivated neutrophils is presented in supplemental figure 5.

### Immunoblotting

Approximately 4 million cells were lysed with Laemmli lysis buffer with phosphatase inhibitor (Thermo Scientific, no. A32957) and HALT protease inhibitor (Thermo Scientific, no. 78425). Proteins of interest were probed with the following antibodies in TBS-T buffer with 5% BSA. Primary antibodies: PYGL (abcam, ab198268), phospho(S15)-PYGL (abcam, ab227043), GLUT1 (Alomone Labs, AGT-041), P-GLUT1 (Millipore Sigma, ABN991), β-actin (Cell Signaling, no. 4967); and secondary antibodies: Goat-anti-rabbit 800 (LI-COR, no. 925-32211), Goat-anti-mouse 680 (LI-COR, no. 925-68070). Membranes were imaged with a Li-Cor Odyssey ClX and quantified using ImageStudio Lite software.

### Glycogen Assay

To extract intracellular glycogen, approximately 3 million cells were aliquoted and pelleted at 300*g* for 2 min. Cells were lysed with ice cold H_2_O and boiled at 100 °C for 15 minutes. Mixture was spun at 1000*g* for 10 min at 4 °C to pellet cell debris. Total glycogen level from the supernatant was measured using the glycogen assay kit (Cayman Chemical, no. 700480) following manufacturer instruction. To quantify glycogen synthesis from extracellular glucose, neutrophils were incubated in media containing U-^13^C-glucose for indicated time. After glycogen extraction as detailed above, glycogen was hydrolyzed with hydrolysis enzyme (Cayman Chemical, no. 700483) at 37 °C for 30 min. Protein was crashed out by addition of 900ul LC-MS grade acetonitrile. Samples were spun at 20,627*g* for 5 min at 4 °C and 850μl of supernatant was transferred to new tube and dried under nitrogen stream. Samples were resuspended in LC-MS water and analyzed by LC-MS.

### Cell viability assay

Cell viability was probed using Cytotox Green (Sartorius 4633) and Annexin V (Sartorius 4641) dyes using an IncuCyte live cell imager, and viability as quantified as the percentage of cells that are negative for both Cytotox Green and Annexin V. To measure cell viability, neutrophils were plated in 96-well culture plate precoated with Cell-Tak (Corning, no. 354240) at 5 × 10^4 cells per well in RPMI media with or without glucose with 0.1% human serum albumin (HSA). Pretreatment of 20μM GPi or DMSO control was added to appropriate wells and the plate was incubated at 37 °C for 1 hour. Cytotox Green and Annexin V dyes were added prior to imaging per manufacturer recommendation. Images were taken every 30–60 min for 24 hours. Images were analyzed using IncuCyte S3 Basic Analysis software.

### Assay for oxidative burst

Oxidative burst was measured by stimulation-induced oxygen consumption rate (OCR) using a XF-96e extracellular flux analyzer (Agilent) following a protocol developed by Seahorse^51^. To attach neutrophils at the bottom of assay plates (Seahorse Bioscience), neutrophils were plated in culture wells precoated with Cell-Tak (Corning, no. 354240) at 4 × 10^4^ cells per well, spun at 200*g* for 1 min with minimal acceleration/deceleration, then incubated for 1 hour at 37 °C. Assays were performed in regular neutrophil culture media (RPMI 1640 medium supplemented with 0.1% human serum albumin) with indicated modifications (glucose withdrawal or GPi treatment). Inhibitors GPi or vehicle control were added just before starting the assay. After several baseline OCR measurements, biochemical stimulant was injected through the injection ports to induce oxidative burst, and the OCR was continuously monitored throughout the course of oxidative burst.

### NET release assay

NET release was quantified by the increase in extracellular DNA after stimulation following a previously developed imaging-based protocol^52^. Briefly, neutrophils were plated at 4 × 10^4 cells per well in a 96-well tissue culture plate precoated with Cell-Tak (Corning, no. 354240) and spun at 200*g* for 1 min with minimal acceleration/deceleration. Cytotox Green Reagent (IncuCyte, no. 4633) was added to culture media at 1:4,000 to stain DNA, and images were captured every 10– 30 min after stimulation using an IncuCyte live cell imager under standard culture conditions (37 °C, 5% CO2). The fluorescent area outside of cells, which indicates NET, was quantified by image analysis using IncuCyte S3 Basic Analysis software. NET release was represented by the ratio of green fluorescence relative to phase area.

### Phagocytosis assay

Neutrophil phagocytosis was quantified by imaging-based analysis using pHrodo-labeled *S. aureus* bioparticles® (Sartorius, no. 4619). The labeled *S. aureus* bioparticles gain fluorescence upon phagocytosis due to the acidic environment of phagosome. Neutrophils were plated at 4 × 10^4^ cells per well in a 96-well tissue culture plate precoated with Cell-Tak (Corning, no. 354240) and spun at 200*g* for 1 min with minimal acceleration/deceleration. Neutrophils were plated in standard culture media with indicated modifications (glucose withdrawal or GPi treatment). pHrodo *S. aureus* bioparticles were added at time 0, and images were captured every 15 min under standard incubation condition (37 °C, 5% CO_2_). Images were analyzed using IncuCyte S3 Basic Analysis software to quantify the fluorescent area normalized to phase area, which indicates phagocytosis.

### Neutrophil migration toward chemoattractant C5a

Migration assays were performed across a C5a gradient through transwell inserts with 3 µm pores (Corning 3415). Isolated primary neutrophils were rested for 30 min in RPMI 1640 medium containing 11mM glucose and 0.1% HSA, then pre-treated for 30 min with experimental condition. Then 2×10^5^ cells were placed into each loading control wells or transwell and migrated towards 2 nM C5a (biotechne, 2037-C5) added to the other side of transwell. Images were taken throughout the migration process and cells migrating to the bottom of each well were quantified using IncuCyte S3 Basic Analysis software. The number of cells migrated after a 2- hour period was corrected for cell loading. Cell loading was corrected for by quantifying the cells in loading control wells and calculating a normalization factor relative to the control group (11mM Glucose, DMSO).

### Fungal control and neutrophil recruitment

To assess neutrophil/fungal interactions, *A. fumigatus* was prepared as following. *A. fumigatus* strain CEA10-RFP (TDGC1.2) was grown on glucose minimal media (GMM) at 37°C in the dark for 3 days prior to the experimental start date to induce asexual conidiation. Following incubation, fungal spores were collected in approximately 25ml of 0.01% Tween 20 solution using an L-spreader and filtered through a 0.4μm mesh filter (Fischer Scientific) into a 50ml conical tube. A 5 ml aliquot was then transferred to a new tube, centrifuged at 800 rpm for 5 min and then resuspended in GMM. Fungal spores were then quantified via hemocytometer and diluted to a final concentration of 4 x 10^3^ cells/ml in GMM. Following dilution, 100 μl of the spore suspension was added to the appropriate number of wells in a 96-well clear, flat bottom plate and left to incubate at 37°C, 5% CO_2_ for 8 hours, or until germlings appeared.

Roughly two hours prior to the end of the spore incubation, primary human neutrophils were collected as described above and diluted to a cell density of 4 x 10^5^ cells/ml in culture medium (RPMI medium with 0.1% human serum albumin). Following dilution, 20μM of GPi or DMSO vehicle control was added to the neutrophil suspension and incubated at 37 °C for 1 hour before the treatment was removed and medium was replaced with regular neutrophil culture medium. After germling development, the fungal culture medium was removed, and 100μl of the neutrophil suspension was added to each well. The plate was then immediately taken to a Nikon Eclipse Ti inverted microscope with a preheated chamber set to 37°C, and initial images, denoted as time 0 (T0), were taken for all wells. All locations imaged in the plate were tracked in the NIS Element software package connected to the microscope. Following image acquisition, the plate was incubated for 4 hours at 37°C, 5% CO_2_. After 4 hours, the plate was removed and imaged at the same locations captured at the start of the experiment (T4). Images were then analyzed for the presence/absence of red hyphal fluorescence between T4 and T0, and for the formation of neutrophil clusters around hyphal cells that were present at the T0 time point. For all conditions, two biological replicates using neutrophils from separate donors with one to three technical replicates for each biological replicate were performed. Statistical differences were assessed via a student’s t-test or a non-parametric Kruskal-Wallis test (GraphPad Prism, v7.0c software).

### Statistics and reproducibility

Statistical analyses were conducted using GraphPad software, with each statistical test listed in the figure legends. Most experiments were repeated independently to ensure reproducibility, as specified in figure legends. Experimental sample sizes used are noted in the figure legends. No statistical method was used to predetermine sample size. In some cases, sample sizes in the same figure across conditions were different mainly due to variation in neutrophil yield from isolation with each blood draw, which can be limited to cover all experimental conditions. Samples are randomly assigned to experimental conditions. The investigators were not blinded to the experiment.

## Acknowledgements

The authors thank Mark Zhang, Mariah Endres, Mitch Howard, and Christina Sheehan for assistance and coordination related to the *in vivo* infusion study. The authors thank Matt Stefely for assistance in figure editing and Alicia Willians for text editing. The authors thank the University of Wisconsin Carbone Cancer Center Flow Cytometry Laboratory, supported by P30 CA014520, for use of its facilities and services This work is supported by NIH grant R35GM147014 (J.F.), R35GM118027 (A.H.), and UW2020 Initiative award AAH8454 provided by the University of Wisconsin - Madison Office of the Vice Chancellor for Research and Graduate Education with funding from the Wisconsin Alumni Research Foundation (J.F., S.M.S, and C.D.F.). Student support was provided in part by T32GM140935 (J.L., N.L.A.), TL1TR002375 (J.L.), and UL1TR002373 (J.L.).

## Author Contributions

E.C.B. and J.F. conceptualized the study and designed the experiments in discussion with other authors. E.C.B., J.L., N.L. A., H.K., S.M.S., C.F., and J.F. contributed to designing or performing the *in vivo* tracing study. E.C.B., X.Q., J.L., and J.A.V. performed neutrophil isolation. E.C.B. and X.Q. performed metabolomics and isotopic tracing experiments in neutrophils. X.Q. performed flow cytometry. J.A.V. performed migration assay with neutrophils. J.L. performed viability assay. A.W. and S.S performed the pathogen control experiments. A.H. contributed to data analysis and interpretation. E.C.B. and J.F. wrote the manuscript with help from all authors.

---

**Supplemental Figure 1.**
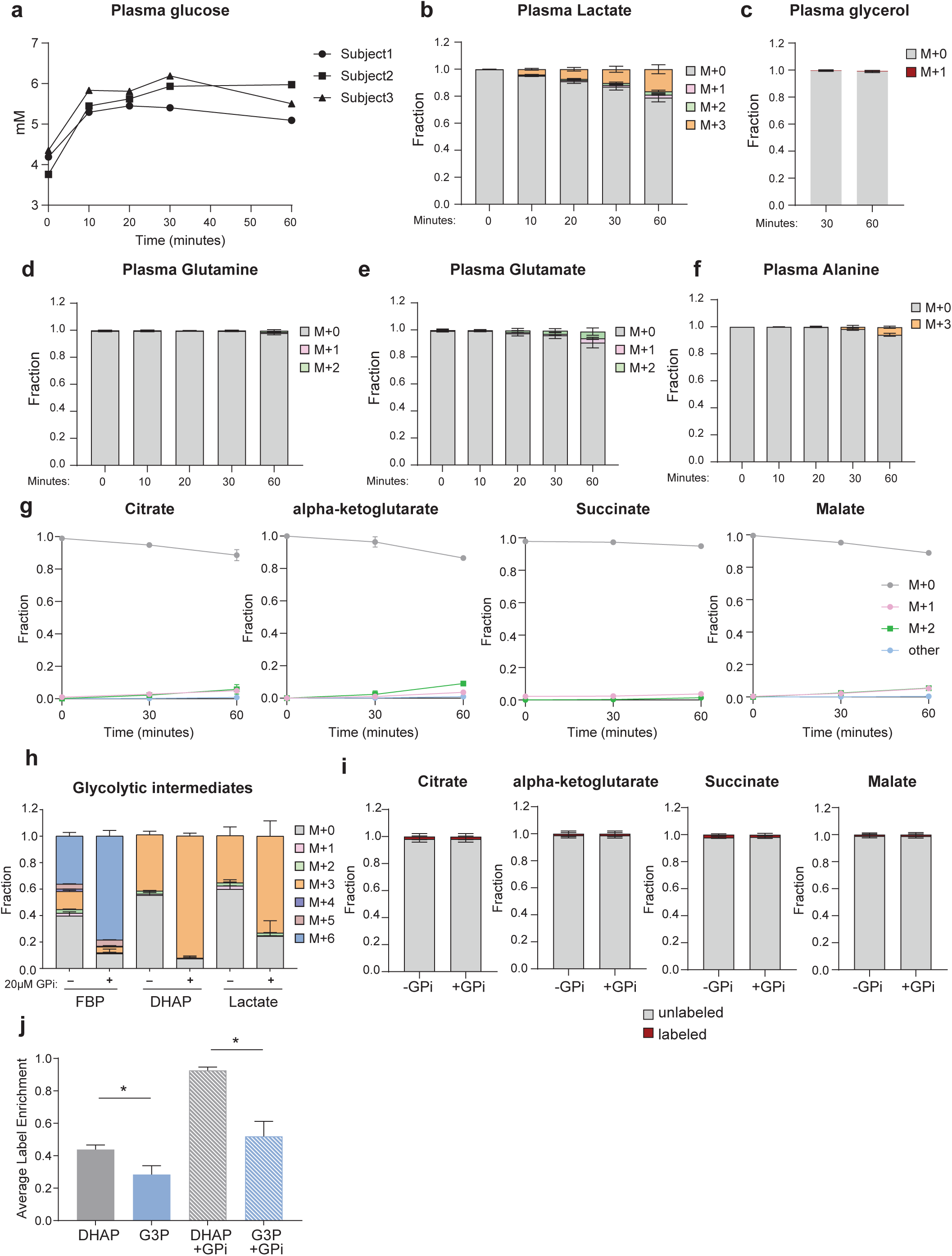
a. Plasma glucose concentration of 3 participants over the time course of U-13C-glucose infusion. b-f. Labeling incorporation into plasma lactate, glycerol, glutamine, glutamate, and alanine from U-13C-glucose over the time course of U-13C-glucose infusion. Results show mean ± SD from n=3 independent participants. g. Labeling incorporation into intracellular TCA cycle metabolites in neutrophils isolated from blood samples 30 or 60 minutes after in vivo U-13C-glucose infusion. Results show mean ± SD from n=3 independent participants. h. Full labeling distribution of glycolytic intermediates in human peripheral blood neutrophils labeled ex vivo in media containing U-13C-glucose for 30min, with or without treatment of GPi (20μM). i. Labeling incorporation into TCA cycle metabolites in peripheral blood neutrophils labeled ex vivo in media containing U-13C-glucose for 30 minutes, with or without GPi (20μM) treatment. j. Average label enrichment of DHAP and glycerol-3-phosphate (G3P) in peripheral blood neutrophils labeled ex vivo in media containing U-13C-glucose for 30min, with or without treatment of GPi (20μM). h-j. Results show mean ± SD from n=3 independent donors.

**Supplemental Figure 2.**
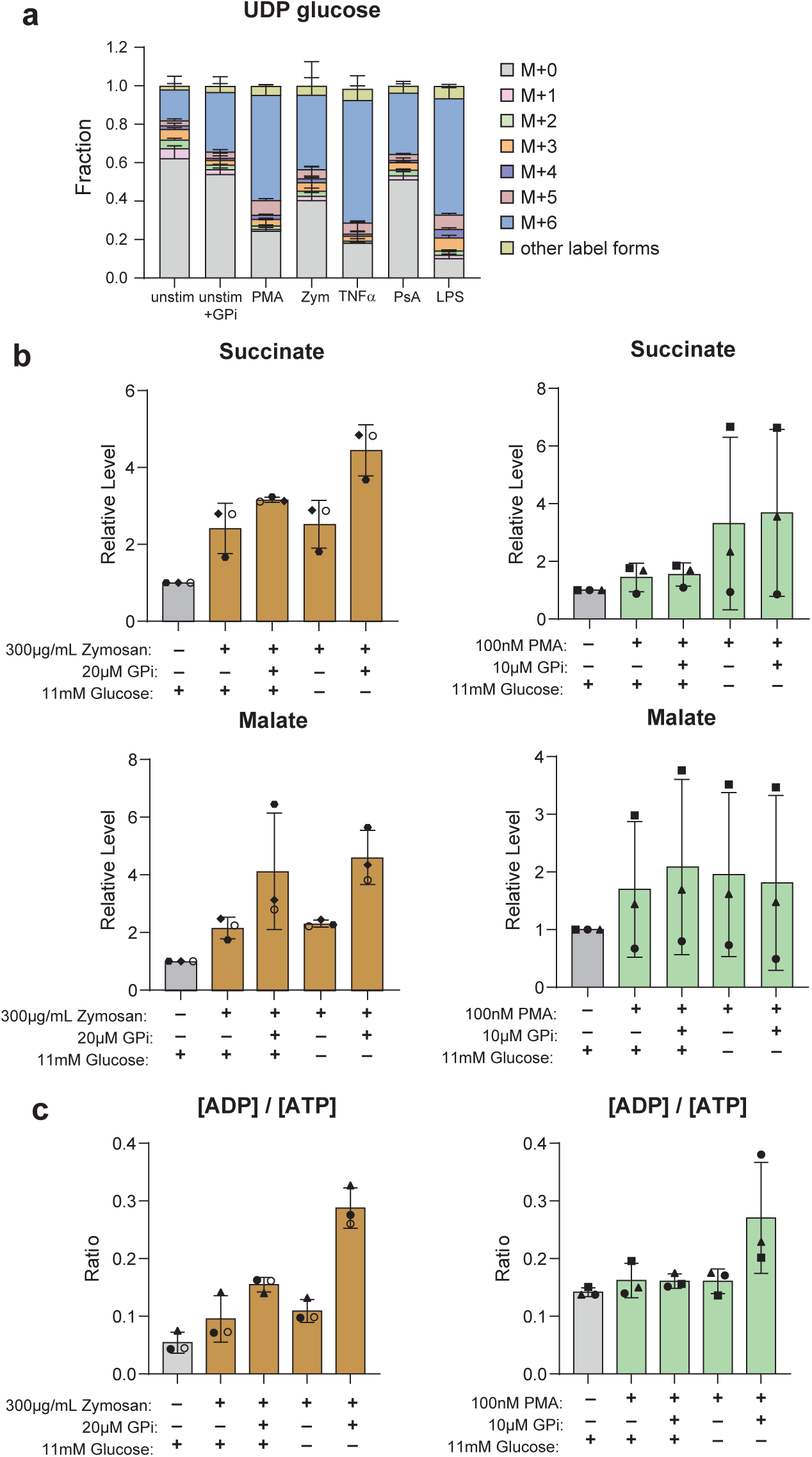
a. Labeling pattern of UDP-glucose from U-13C-glucose in unstimulated neutrophils with (n=3 donors) or without (n=5 donors) treatment of 20μM GPi, or neutrophils stimulated with 100nM PMA (n=4 donors), 300μg/mL zymosan A (n=6 donors), 100ng/mL TNFα (n=4 donors), or heat inactivated P. aeruginosa (HI PsA, 10:1 bacteria: neutrophil) (n=3 donors) for 30 minutes or stimulated with 1ug/mL LPS for 4 hours (n=3 donors). Results show mean ± SD. b-c. Relative levels of TCA cycle intermediates (b) and ADP/ATP ratio (c) in unstimulated neutrophils or neutrophils stimulated with zymosan A or PMA cultured in media with indicated nutrient perturbation (with or without glucose, with or without GPi treatment). Bars and error bars show mean ± SD from n=3 independent donors, as represented by individual dots.

**Supplemental Figure 3.**
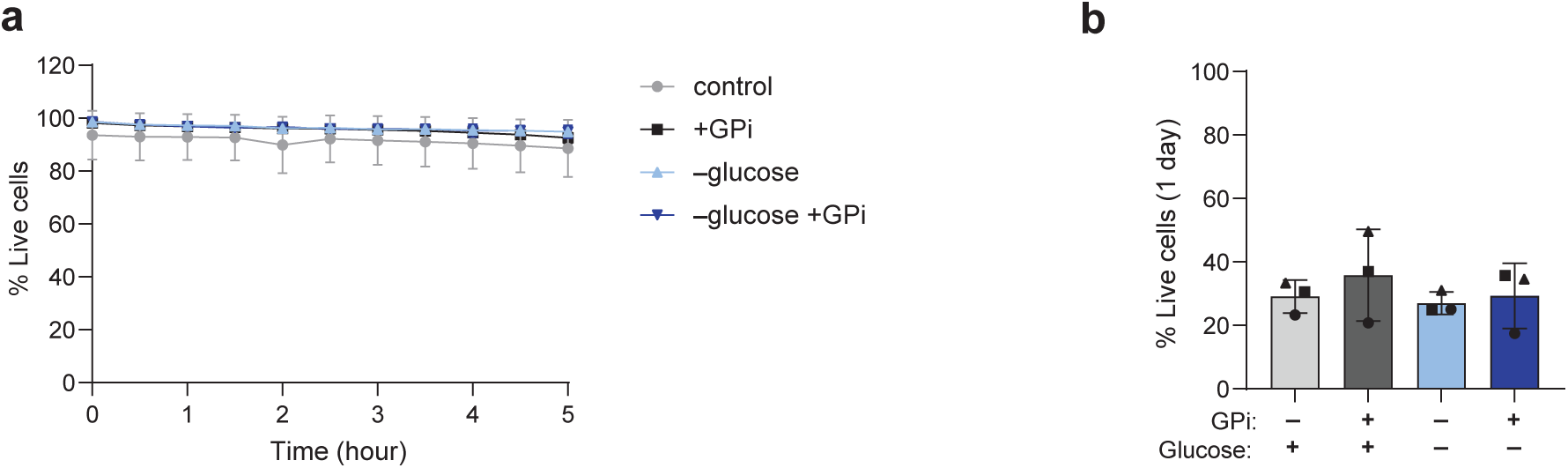
a. Viability of unstimulated neutrophils over a course of 5h when cultured in indicated media condition (with or without glucose, with or without 20μM GPi treatment). Control is standard RPMI media-based culture condition. ‘Live cells’ are the cells that are negative of both Cytotoxic and annexin V stain. Mean ± SD from n=3 independent donors. b. Percent of live cells after unstimulated neutrophils were cultured in indicated media condition for 1 day (23-24h). Bars and error bars show mean ± SD from n=3 independent donors, as represented by different markers.

**Supplemental Figure 4.**
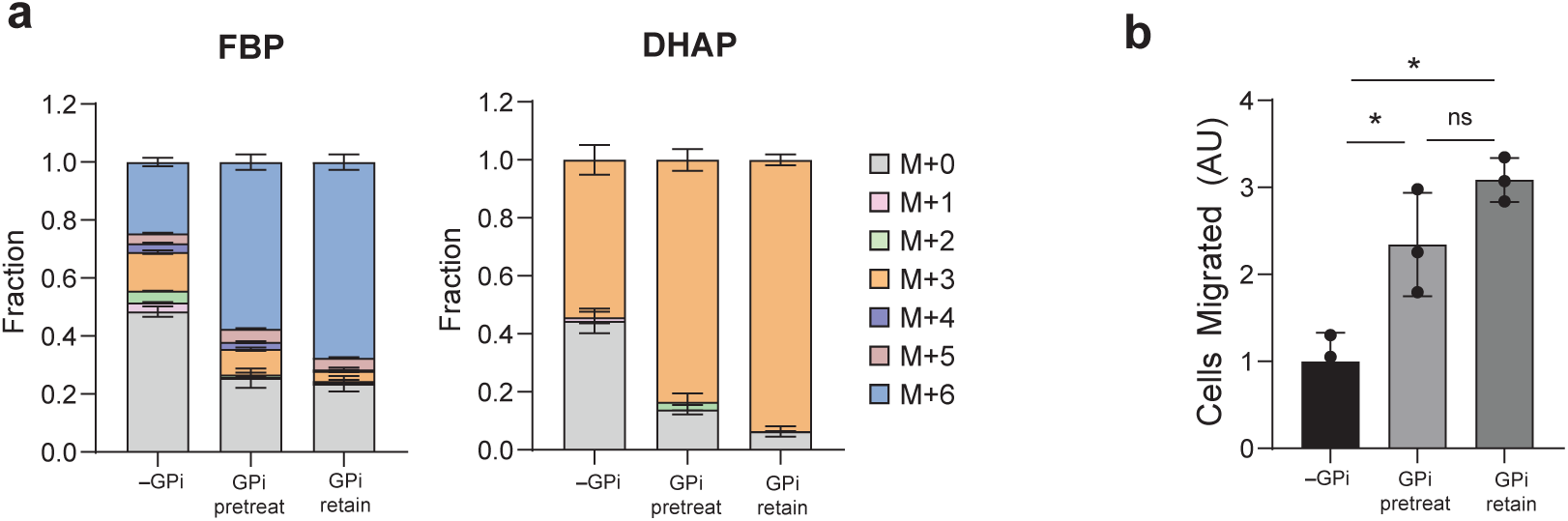
a. Labeling pattern of intracellular glycolytic intermediates following 4-hour incubation in U-13C-glucose media in neutrophils that are untreated, treated with 20μM GPi throughout the duration of labeling, or pre-treated with 20μM GPi for 1hr prior to labeling in media containing no GPi. Bars and error bars show mean ± SD from n=3 technical replicates. b. Migration of neutrophils through transwell towards 2nM C5a. Cells were untreated (-GPi), pretreated with GPi for 30 minutes prior to addition to transwell without GPi for migration assay (GPi pretreat), or treated with 20uM GPi throughout the duration of migration assay (GPi retain). Bars and error bars show mean ± SD from n=3 technical replicates (indicated by individual dots). Statistical analysis was performed by unpaired t-test.

**Supplemental Figure 5.**
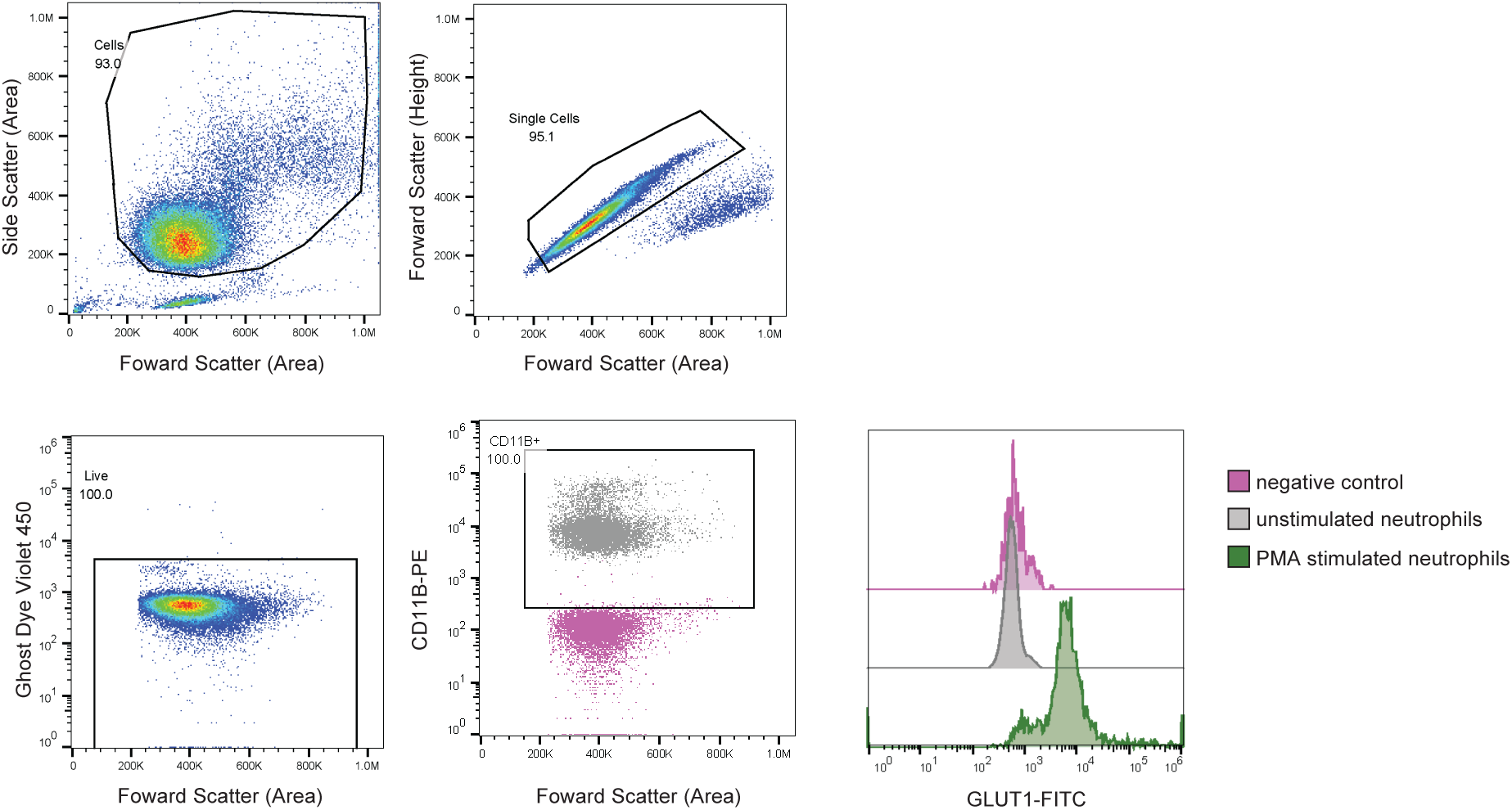
Gating strategy for interrogation of GLUT1 on the surface of unstimulated neutrophils. Isolated cells were stained with Ghost Dye Violet 450, CD11B-PE, and GLUT1-FITC.

